# Systematic dissection of Cas12a-mediated precision genome editing defines design principles for genome-scale variant engineering

**DOI:** 10.64898/2026.06.26.734799

**Authors:** Antoine Delhaye, Vlad Ştefan Băţăgui, Jana Nysten, Daan Troubleyn, Sibylle C. Vonesch

**Author notes:** these authors contributed equally (Antoine Delhaye, Vlad Ştefan Băţăgui).

## Abstract

Cas9 precision editing is increasingly predictable because guide, donor and target-context effects have been systematically characterized. Extending this framework to other nucleases is essential for installing variants outside convenient Cas9 target space. Cas12a provides a T-rich protospacer-adjacent motif (PAM) alternative, but determinants of efficient donor-templated Cas12a editing remain poorly defined. Here, we systematically dissected Cas12a precision editing in *Saccharomyces cerevisiae* across nuclease, direct repeat, expression, crRNA, donor, genomic context and time-course variables. Reporter and amplicon-sequencing assays showed that cleavage activity alone did not predict precise editing. Highly active configurations often reduced viability or lost edited alleles over time, whereas attenuated configurations better preserved programmed edits. Enhanced AsCas12a edited rapidly and tolerated shorter crRNAs, resulting in a narrower editing window, while an attenuated FnCas12a configuration edited more slowly but maintained higher viability and better distal-edit recovery. Alternative repair outcomes were rare, target-dependent, and further suppressed by LexA-FHA donor recruitment. To define design parameters at scale, we established a pooled Cas12a platform with 530 barcoded edit cassettes and recovered programmed edits for 70.2% of designs. Successful editing was reduced with TTTG PAMs, a C upstream of the PAM and at distal edit positions. Excluding these features increased the edited fraction to 85.4% and adding high predicted cleavage scores further elevated it to 91.4%. Applied retrospectively, these criteria also identified poorly edited loci in the targeted panels. Together, these data define design principles for Cas12a-mediated precision editing and establish a scalable platform for genome-scale pooled variant engineering and phenotyping in yeast.

## Background

Clustered regularly interspaced short palindromic repeats (CRISPR) and CRISPR-associated (Cas) proteins have transformed research, biotechnology and medicine by enabling programmable nucleic acid recognition and cleavage [1]. In genome editing applications, CRISPR nucleases generate a double-strand break (DSB) at a defined genomic locus, after which the lesion is repaired by endogenous DNA repair pathways [2]. Non-homologous end joining (NHEJ) can eventually produce small insertions or deletions and is therefore widely exploited for gene disruption [3]. In contrast, homology-directed repair (HDR) can install precise sequence changes when a donor template carrying the desired edit flanked by homology to the targeted locus is supplied [4]. Because many applications require defined alleles rather than disruptive mutations, improving the efficiency and predictability of HDR remains a central challenge in genome editing.

Although Cas9 remains the most widely used nuclease, Cas12a offers several features that make it an attractive complementary platform for precision genome engineering. Unlike Cas9, Cas12a uses a single crRNA and does not require a tracrRNA, and it recognizes targets adjacent to a T-rich PAM [5], thereby expanding the editable sequence space. The crRNA contains a direct repeat (DR) that folds into a stem-loop/pseudoknot structure and is structurally highly conserved across orthologs [5,6]. The DR is essential for autonomous pre-crRNA processing [7], as well as for nuclease loading and activity [5]. Following PAM recognition, Cas12a initiates target binding through PAM-proximal pairing and forms a directional R-loop that extends into an approximately 20-bp crRNA-target-strand hybrid [7,8]. Because R-loop propagation remains reversible until late in target interrogation, Cas12a is sensitive to mismatches along much of the target sequence [9], providing a mechanistic basis for its high specificity [10]. This mismatch sensitivity may also be advantageous for precision editing without dedicated PAM-blocking mutations, because the desired edit itself can reduce re-cleavage of the edited allele. Cas12a then cleaves DNA distal to the PAM, producing staggered double-strand breaks with 5′ overhangs [5] rather than the predominantly blunt ends generated by Cas9 [1].

Despite this detailed mechanistic understanding, Cas12a has been characterized mainly through target-cleavage and indel-formation assays [11,12], whereas the determinants of donor-templated HDR remain less well defined. For Cas9, several principles governing precision editing efficiency have been established, including the importance of guide activity, donor design, target sequence and genomic context and the distance between the intended edit and the cleavage site [13–16]. Equivalent rules for Cas12a-mediated HDR remain less well-defined. Work in mammalian cells suggests that placing edits around positions 12-16 within the protospacer can maximize HDR efficiencies [14], but the generality of this observation across targets, nucleases and expression architectures remains unclear. More broadly, the extent to which factors like Cas12a ortholog choice, nuclease expression level, crRNA architecture and expression, target context and donor design shape HDR efficiency, competing repair outcomes and editing viability remains poorly understood.

The yeast *Saccharomyces cerevisiae* is a powerful model for addressing these questions. Its efficient homologous recombination machinery makes it unusually tractable for systematically dissecting how HDR outcomes are shaped by the editing machinery and cassette design parameters. Several Cas12a orthologs have already been shown to function in yeast [17–19], and recent studies have further optimized individual Cas12a systems [20] and PAM-relaxed variants [21] for yeast engineering. However, these studies have largely focused on activity at selected loci or on overall editing performance rather than on the determinants of precise HDR itself, limiting both mechanistic understanding and rational design.

Here, we used yeast to systematically define the determinants of Cas12a-mediated precision editing. We first benchmarked five Cas12a orthologs across nuclease expression levels, direct repeat configurations and crRNA promoters using complementary reporter assays that distinguish overall editing from HDR repair. We then focused on the two best-performing systems to determine how target context, crRNA length, and mutation position relative to the PAM influence HDR efficiency, alternative repair outcomes and viability at reporter and endogenous loci. Finally, we established a pooled platform for Cas12a-mediated variant installation and combined it with low-coverage whole-genome sequencing to resolve editing outcomes across hundreds of endogenous targets. This genome-scale analysis identified sequence and cassette design features associated with editing success. Together, these analyses define key determinants of Cas12a precision editing and establish a pooled Cas12a platform that broadens the target space for genome-scale precision editing screens in *S. cerevisiae*.

## Results

### Comparative benchmarking of five Cas12a orthologs and various expression systems for cleavage and precision editing

To benchmark different Cas12a orthologs for HDR-mediated precision editing in yeast, we selected a phylogenetically diverse panel of variants with prior evidence of activity in yeast and/or mammalian systems. As mammalian codon-optimized versions of FnCas12a, AsCas12a, LbCas12a, and MbCas12a have been reported to function in yeast [19], we included these variants together with yeast codon-optimized versions of LbCas12a and FnCas12a [18]. In addition, we included enhanced AsCas12a (enAsCas12a) due to its increased activity in mammalian cells [22]. To ensure robust and comparable constitutive expression, all Cas12a constructs were expressed from the *TEF1* promoter and crRNAs from the *SNR52* promoter, a combination that outperformed other expression architectures for Cas9 [16]. In addition to each cognate DR, we tested the AsCas12a DR as a common non-cognate repeat because it has been reported to be tolerated by several Cas12a orthologs [5,19]. This allowed us to test cross-ortholog compatibility and assess whether DR exchange could tune editing activity. Editing efficiency was benchmarked by introducing an HDR donor-encoded premature termination codon (PTC) in a genomically integrated sfGFP (Fig. 1A). To compare editing dynamics, total editing was quantified at different timepoints by flow cytometry as the fraction of GFP-negative cells, capturing all repair outcomes that disrupt fluorescence.

**Figure 1.**
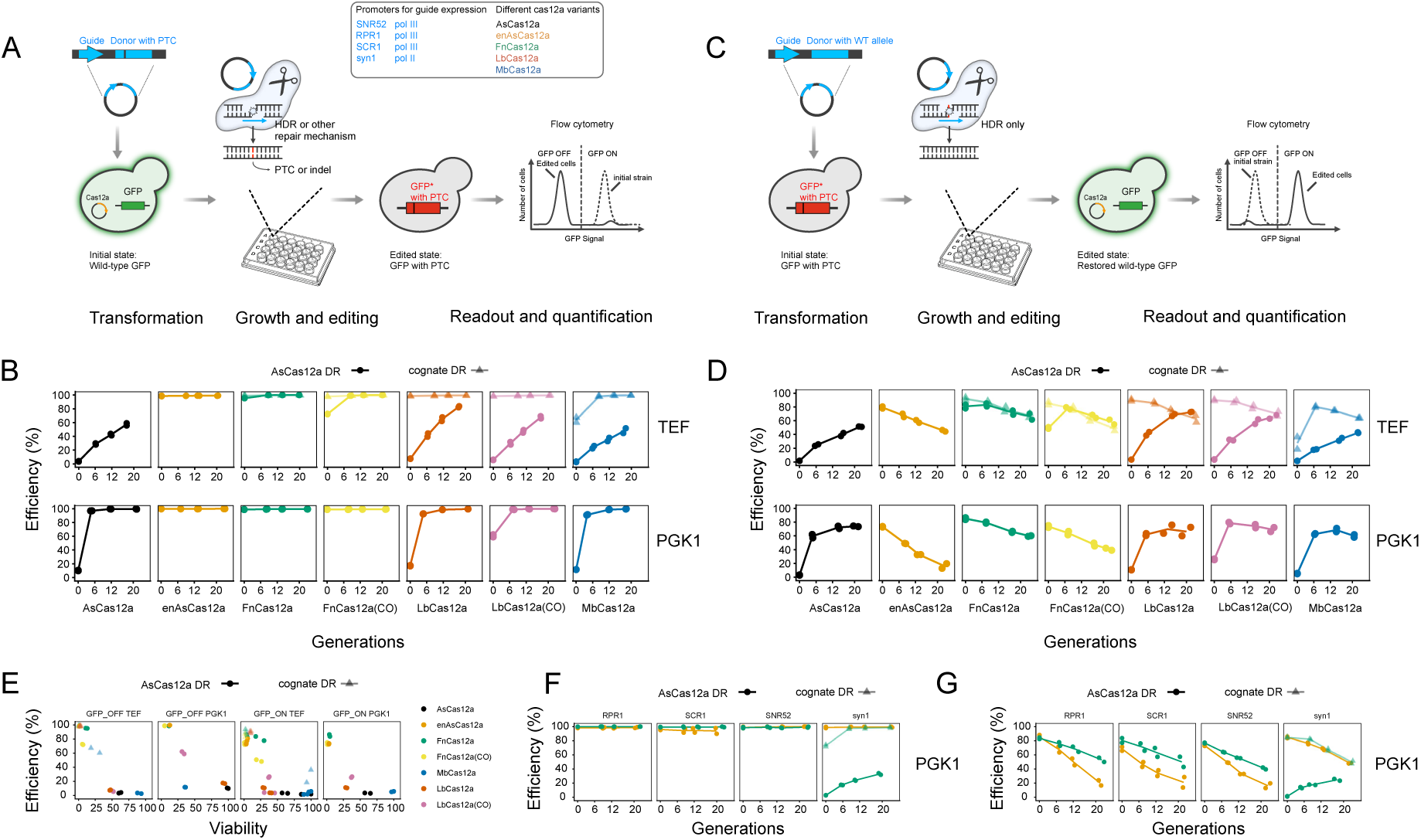
Comparative benchmarking of Cas12a orthologs and expression architectures using complementary GFP reporter assays. (A) GFP-loss assay. Yeast carrying a genomic sfGFP reporter were transformed with plasmids encoding the indicated Cas12a variant, a GFP-targeting crRNA, and an HDR donor designed to introduce a premature termination codon (PTC) in GFP. Cas12a and crRNAs were expressed from indicated promoters. Editing was quantified by flow cytometry as the fraction of GFP-negative cells (n = 25,000 cells), capturing donor-templated HDR and other GFP-disrupting repair outcomes. Measurements were taken after 2 days of plate incubation (generation 0) and after 6, 12, and 20 additional generations of editing in liquid culture. Two plated replicates per transformation were propagated independently. (B) GFP-loss kinetics across Cas12a orthologs expressed from p*TEF1* (top) or p*PGK1* (bottom), with crRNAs expressed from p*SNR52*. (C) GFP HDR assay. A GFP allele carrying the PTC introduced in the GFP-loss assay was targeted with a crRNA and HDR donor designed to restore the wild-type sequence. GFP-positive cells therefore arise only after donor-templated HDR, allowing HDR efficiency to be quantified as the fraction of GFP-positive cells. All other details are as in (A). (D) HDR kinetics across Cas12a orthologs. (E) Efficiency-viability tradeoff. Facets show editing efficiency and relative viability for each Cas12a ortholog in the GFP-loss and GFP-restoration assays, with p*TEF1*- or p*PGK1*-driven Cas12a expression and *pSNR52*-driven crRNA expression. Editing efficiency data are the same as in B and D. Viability was calculated from colony counts after plating normalized within each experiment (see Methods). (F) Effect of crRNA expression on GFP-loss kinetics using p*PGK1*-driven enAsCas12a (gold) or FnCas12a with As DR (green). (G) Effect of crRNA expression on HDR kinetics. In B and D-G, colors indicate Cas12a orthologs, shapes indicate direct repeats (circle = As DR, triangle = cognate DR), symbols show plated replicates and lines indicate replicate means. FnCas12a(CO) and LbCas12a(CO) refer to the yeast codon-optimized constructs.

Most orthologs reached maximal or near-maximal editing with their cognate DR except for AsCas12a, which reached only 60% efficiency (Fig. 1B, top panel, Supplementary Table S1). MbCas12a exhibited slower kinetics than FnCas12a, LbCas12a, and enAsCas12a, reaching 100% editing after six additional generations. In contrast, only FnCas12a retained full activity with the AsCas12a DR, whereas LbCas12a and MbCas12a showed reduced efficiency and delayed kinetics (Fig. 1B, top panel). Comparison of human- and yeast-optimized coding sequences showed little overall effect of codon optimization, although yeast-optimized FnCas12a edited more slowly than its human-optimized counterpart (Fig. 1B, top panel). Although the GFP-loss assay provides a rapid readout of editing activity, it does not distinguish precise HDR from other fluorescence-disrupting outcomes, including NHEJ-mediated indels or larger deletions generated by microhomology-mediated end joining (MMEJ) or single-strand annealing (SSA). We therefore used a complementary GFP restoration assay to quantify HDR specifically. In this assay, a genomically integrated GFP allele carrying the previously introduced PTC is restored to the wild-type sequence, such that GFP-positive cells arise only through HDR-mediated repair (Fig. 1C). Using the AsCas12a DR, the weaker or slower editors in the GFP-loss assay, namely AsCas12a, LbCas12a, and MbCas12a, showed a progressive accumulation of HDR products over time (Fig. 1D, top panel; Supplementary Table S1). In contrast, enAsCas12a and FnCas12a showed high initial HDR frequencies that decreased at later time points (Fig. 1D, top panel), with FnCas12a reaching higher HDR levels overall and exhibiting a more modest decline. A similar temporal decline was observed for most orthologs when paired with their cognate DRs. Because all orthologs reached maximal or near-maximal editing with their cognate DR, we next asked whether changing Cas12a expression could enhance activity with the non-cognate DR. Expression from p*PGK1*, a glycolytic promoter widely used for constitutive expression in yeast, substantially improved editing by LbCas12a and MbCas12a (Fig. 1B, bottom panel). HDR efficiency also increased, although more modestly and most clearly at early time points (Fig. 1D, bottom panel). With p*TEF1*-driven expression, overall editing and HDR efficiency were strongly correlated for the weaker editors, including LbCas12a, MbCas12a, and AsCas12a. In contrast, p*PGK1*-driven expression primarily increased overall editing (Supplementary Fig. 1A). Together, these data indicate that the most active Cas12a-DR combinations are not necessarily the most stable over time and highlight DR compatibility, temporal behavior and expression as key variables when comparing Cas12a orthologs for precision editing.

A further relevant parameter for precision editing, particularly in large-scale screens, is the balance between nuclease activity and cell viability. Consistent with cleavage-associated toxicity, less active Cas12a-DR combinations generally showed higher relative viability after editing (Fig. 1E). Among the strongest editors, FnCas12a (As DR) yielded the highest colony counts, indicating a favorable balance between activity and viability (Fig. 1E; Supplementary Table S1). Growth assays further showed that this fitness cost was mainly cleavage dependent, as strains expressing non-targeting crRNAs grew similarly to the empty-plasmid control (Supplementary Fig. 1B-D). In contrast, GFP-targeting crRNAs reduced growth for several variants, particularly when paired with cognate DRs. Constitutive Cas12a and crRNA expression therefore imposed little basal burden, with fitness defects mainly reflecting active DNA cleavage and repair at the target locus.

Finally, we asked whether crRNA promoter choice further modulated editing efficiency and precision. To this end, we tested a panel of crRNA promoters for two strong editors enAsCas12a and FnCas12a (As DR), both expressed from the *PGK1* promoter, a condition under which both systems showed maximal editing at the earliest measured timepoint. In addition to p*SNR52* tested above, we included the strong yeast Pol III promoter p*RPR1*, as well as p*SCR1*, a weaker Pol III promoter with distinct promoter architecture and reported CRISPR compatibility in yeast [23]. We also tested syn1, a strong fully synthetic yeast Pol II promoter (Supplementary Fig. 2) [24]. Because Pol II-driven Cas12a crRNA expression requires transcription of a DR-spacer-DR precursor followed by Cas12a-mediated processing [25], the syn1 design tested both promoter strength and pre-crRNA maturation by each Cas12a-DR combination (Supplementary Fig. 2). Across the Pol III promoters tested, both enAsCas12a and FnCas12a (As DR) supported maximal or near-maximal total editing (Fig 1F). As in the previous experiment, HDR decreased over time, and this reduction was more pronounced for enAsCas12a (Fig. 1G). In contrast, syn1-driven crRNA expression strongly reduced FnCas12a editing when combined with the non-cognate AsCas12a DR, whereas this defect was alleviated when FnCas12a was paired with its cognate DR (Fig. 1F, G). Because the Pol II architecture requires crRNA maturation, these results suggest that non-cognate DRs can remain compatible with ribonucleoprotein (RNP) assembly and cleavage yet be less efficient during precursor processing.

Based on these results, we selected enAsCas12a and FnCas12a (As DR) for further characterization. We used p*TEF1* for Cas12a expression in all subsequent experiments, as enAsCas12a showed a less pronounced decline in HDR under this condition (Fig. 1D), while FnCas12a (As DR) retained higher post-editing viability than under p*PGK1* expression (Fig. 1E, Supplementary Table S1). For crRNA expression, we selected the Pol III promoters *pSNR52* and p*RPR1*, which provided simpler crRNA expression architectures than the Pol II design and supported maximal editing for both nucleases. We retained *pRPR1* alongside *pSNR52* because HDR decline was less pronounced over time. Together, these nuclease and promoter combinations provided a useful range of editing regimes, from rapid high-activity editing with stronger HDR decline to more attenuated configurations with greater temporal stability.

### Cas12a editing across a GFP crRNA panel varies with crRNA, ortholog, and donor-dependent parameters

Because editing outcomes can be strongly target-dependent, we next evaluated enAsCas12a and FnCas12a (As DR) across 12 crRNA/donor pairs designed to introduce PTCs in a genomically-encoded sfGFP reporter by single nucleotide substitutions (Fig. 2A, Supplementary Table S2). The panel captured features expected to influence cleavage and HDR efficiency, including crRNA GC content and the distance between the desired edit and the PAM (Supplementary Table S2). A non-targeting crRNA with minimal homology to GFP or the yeast genome, paired with a donor, served as a negative control. Both nucleases supported high levels of editing for most crRNAs (Fig. 2B, Supplementary Table S3), but editing kinetics varied across targets and could be grouped qualitatively into distinct kinetic behaviors, including progressive increases, stable high-editing trajectories, transient peaks followed by decline, and persistently low-editing crRNAs (Fig. 2C, Supplementary Fig. 3A). EnAsCas12a generally achieved maximal or near-maximal editing after 6 generations of liquid outgrowth under all tested conditions, except for two crRNAs that were poorly functional or associated with low viability and potential escaper enrichment (Fig. 2B, C, Supplementary Fig. 3A, B). FnCas12a (As DR) also achieved high final editing levels at most targets but typically with slower kinetics, resulting in lower average editing at intermediate time points and greater sensitivity to crRNA promoter choice, with p*RPR1* performing less well than p*SNR52* (Fig. 2C, Supplementary Fig 3A, B). Target-dependent effects were also evident, with some crRNAs performing better with FnCas12a (As DR), others with enAsCas12a, and promoter effects varying across crRNAs (Fig. 2B, Supplementary Table S3). These target-dependent differences were most apparent at early time points, before many crRNAs converged on 90-100% editing, highlighting editing kinetics as an additional dimension for evaluating crRNA efficacy. The panel included matched crRNA-donor comparisons in which the same crRNA was paired with different donors to introduce distinct edits. These comparisons showed that donor design can strongly affect editing efficiency and viability. For enAsCas12a, donors placing the edit more distally from the PAM consistently performed worse than the matched PAM-proximal donors, except when both edits were within 5 nucleotides (nt) of the PAM (Supplementary Fig. 4). In contrast, FnCas12a (As DR) showed more similar editing efficiency and viability across matched donor pairs (Supplementary Fig. 4A, B). Thus, beyond crRNA activity, donor-dependent parameters are major determinants of productive editing and cell survival, and their effects can differ between nucleases.

**Figure 2.**
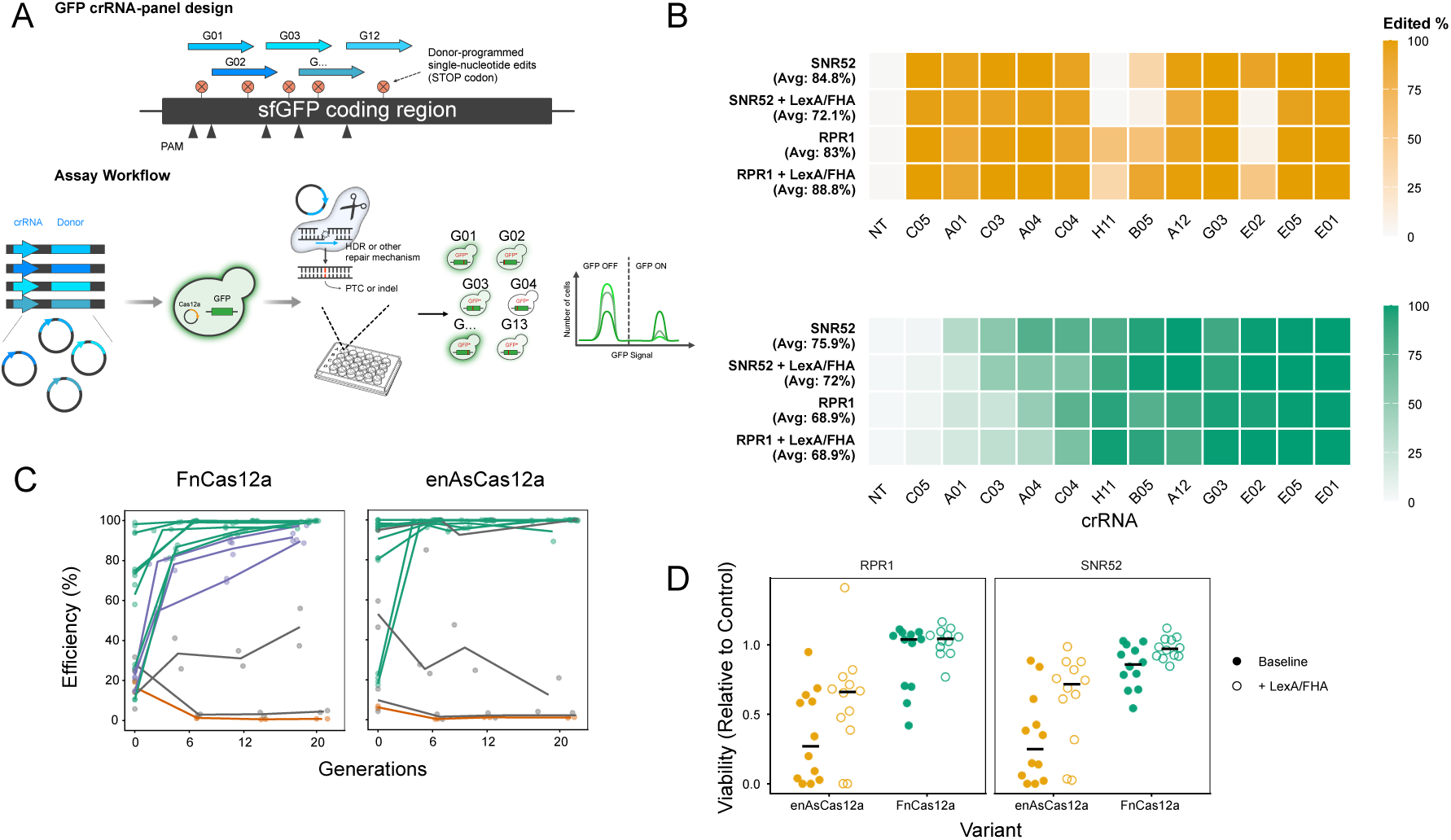
Editing across a GFP stop-codon panel reveals effects of crRNA promoter and donor recruitment. (A) GFP crRNA panel screen. Yeast carrying a genomic sfGFP reporter were transformed with plasmids encoding p*TEF1*-driven enAsCas12a or FnCas12a (As DR), a GFP-targeting crRNA, and an HDR donor designed to introduce a PTC in GFP. The panel contained 12 GFP-targeting crRNA-donor pairs spanning different target positions, PAM sequences, target-strand orientations, and PAM-to-edit distances, and one non-targeting control. crRNAs were expressed from indicated promoters and editing performed with or without LexA-FHA-mediated donor recruitment. Editing was quantified by flow cytometry as the fraction of GFP-negative cells (n = 25,000 cells) after plating and after an additional 6, 12, and 20 generations of editing in liquid culture. Two plated replicates per transformation were propagated independently. (B) Editing efficiency across the GFP crRNA panel after 6 generations of editing in liquid culture. Heatmaps show average GFP loss for enAsCas12a (top, gold) and FnCas12a (As DR) (bottom, green) across editing conditions. Columns represent the non-targeting control (A09) followed by GFP-targeting crRNA-donors labelled with well id and ordered by mean FnCas12a editing efficiencies. Rows indicate editing conditions, fill values correspond to average efficiency across plated replicates, and the average efficiency across targeting crRNAs per condition is shown on the left. (C) GFP-loss kinetics across the GFP crRNA panel for FnCas12a (As DR) (left) and enAsCas12a (right), with p*SNR52*-driven crRNA expression without donor recruitment. Points indicate replicates and lines means for each target. Colors indicate kinetic classes assigned separately for each nuclease: green = high (>90% editing at 6 generations), grey = low (never exceeds 50% mean editing), purple = intermediate, orange = nontargeting crRNA. (D) Relative viability across the GFP crRNA panel. Panels show p*RPR1*- or p*SNR52*-driven crRNA expression, with points colored by Cas12a variant and fill values denoting donor-recruitment condition. Relative viability represents colony counts normalized to the non-targeting control in each experiment.

We next asked whether editing with poorly performing crRNAs could be enhanced by donor recruitment. To this end, we repeated experiments with the LexA-FHA system, which recruits repair templates to DNA double-strand breaks and improves both HDR efficiency and cell survival during Cas9-mediated editing in yeast [15,26]. For both nucleases, donor recruitment had modest and target-dependent effects, improving editing at some targets while having little or even adverse effects at others (Fig. 2B, Supplementary Fig. 3A, B). In contrast, viability increased substantially across crRNAs (Fig. 2D, Supplementary Table S3). Under the conditions tested here, the main benefit of donor recruitment therefore appears to be partial uncoupling of the editing-viability trade-off, enabling better survival with highly active nuclease-crRNA combinations (Supplementary Fig. 3C). Because this assay measures total editing rather than HDR specifically, however, potential effects of donor recruitment on Cas12a-mediated precise repair remain to be determined.

### Cas12a precision editing depends on crRNA length and edit position relative to the PAM

All experiments above used 23 nt crRNAs. However, Cas12a can cleave with a crRNA-target complementarity as short as 17 nt [5]. We therefore asked whether crRNA length quantitatively affects HDR efficiency and repair outcomes. We selected a crRNA/donor pair that edited efficiently with enAsCas12a and showed intermediate activity with FnCas12a (As DR) and shortened the crRNA in single nucleotide increments from 23 to 16 nt while keeping the donor template constant. Editing was quantified by amplicon sequencing to directly measure HDR and alternative repair outcomes. enAsCas12a was relatively tolerant of crRNA shortening, maintaining high HDR efficiencies with crRNAs from 23 to 17 nt (Fig. 3B, Supplementary Table S4). In contrast, FnCas12a (As DR) showed a sharp drop in HDR below 19 nt (Fig. 3B). Among crRNA lengths that supported efficient editing, colony counts were highest with longer crRNAs for enAsCas12a (21-23 nt; Fig. 3D) and with length 19 and 22 nt for FnCas12a (Fig. 3E), with an apparent optimum around 22 nt for both nucleases. Thus, crRNA length affects both HDR efficiency and cell survival, although the optimum may be target dependent. Across the crRNA length series, non-HDR outcomes remained infrequent for both nucleases (Supplementary Fig. 5A, C), indicating that crRNA shortening primarily shifted outcomes between precise HDR and retention of the unedited allele rather than broadly redirecting repair toward indel formation.

**Figure 3.**
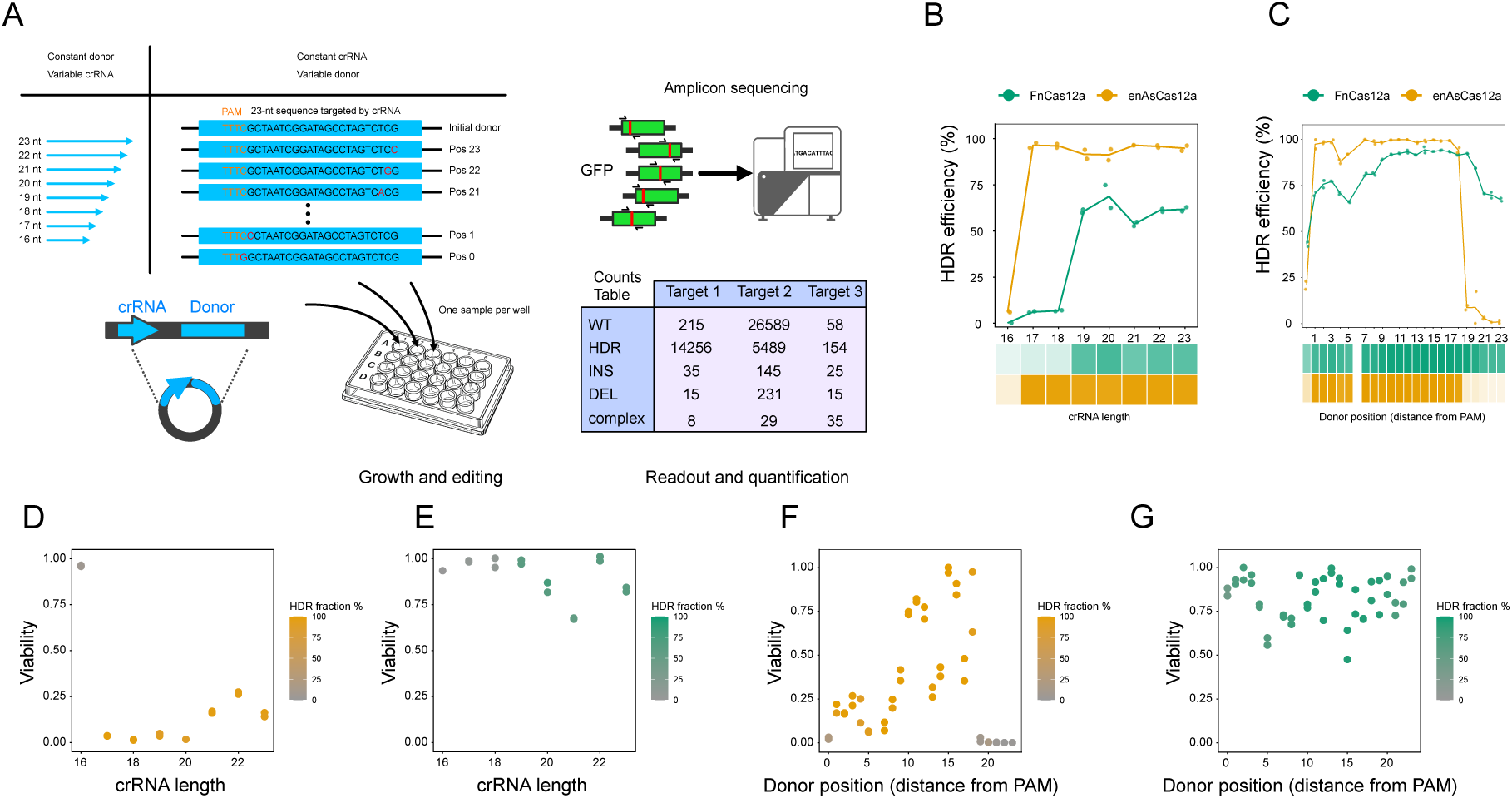
crRNA length and edit position influence Cas12a-mediated HDR efficiency and viability. (A) Experimental design. A single GFP target (A04) was used to create either a crRNA-length series, in which the targeting sequence was truncated from 23 nt to 16 nt while keeping the donor constant, or an edit-position series, in which a single donor-programmed substitution was moved across positions 0 to 23 relative to the PAM. HDR efficiency was quantified by amplicon sequencing after plating and after an additional 6, 12, and 20 generations of editing in liquid culture. Two plated replicates per transformation were propagated independently. (B) Effect of crRNA length on HDR efficiency. Points show replicate HDR measurements across crRNA lengths and lines indicate means. The selected comparison timepoint is 6 generations for enAsCas12a and 12 generations for FnCas12a, corresponding to the most informative dynamic range for each nuclease. The heatmap represents average HDR efficiency for each crRNA length and nuclease. (C) Effect of edit position relative to the PAM on HDR efficiency. Position 0 corresponds to V in a TTTV PAM and position 23 to the distal end of the protospacer. Points and lines are shown as in (B), using the same nuclease-specific comparison timepoints. The heatmap represents average HDR efficiency for each edit position and nuclease. Effect of crRNA length (D) or edit position (E) on relative viability for enAsCas12a. Point color indicates HDR efficiency at 6 generations. Effect of crRNA length (E) or edit position (G) on relative viability for FnCas12a with the As DR. Point color indicates HDR efficiency at 12 generations.

We next examined the effect of edit position, a parameter known to influence precision editing in Cas9 systems, where efficiency generally decreases as the edit is placed further from the PAM or cleavage site [4,14]. We constructed a donor series in which a single programmed base substitution was systematically moved from position 0 to position 23, with position 0 corresponding to the V of the TTTV PAM and position 23 to the distal end of the crRNA. At each position, the donor introduced a single-base transversion while all other donor features were kept constant (Fig. 3A). Position 6 was excluded from the analysis because the corresponding donor plasmid contained an unintended error. HDR remained high across most of the tested range for both nucleases but showed clear position-dependence (Fig. 3C, Supplementary Table S4). enAsCas12a exhibited two windows of high HDR efficiency, spanning positions 1-3 and 7-18. FnCas12a (As DR) showed highest HDR for edits between positions 9 and 19, with a second local peak at positions 1-3. Both nucleases showed a dip in HDR after position 3, followed by recovery. At more distal positions, HDR efficiency declined gradually for FnCas12a (As DR) and more sharply for enAsCas12a, falling below 15% at position 19 and approaching zero for edits placed further downstream. One possible interpretation is that enAsCas12a retains efficient target cleavage with shorter crRNA-target complementarity, such that edits placed beyond this window fail to sufficiently disrupt re-cleavage after HDR editing. Accordingly, enAsCas12a colony counts decreased sharply when edits were placed beyond position 18 (Fig. 3F). Reduced viability was also observed for some PAM-proximal positions, indicating that these mismatches did not fully protect edited alleles from cleavage, consistent with reports that engineered enAsCas12a can retain activity on PAM-proximal mismatched substrates that strongly impair wild-type AsCas12a [27]. In contrast, FnCas12a (As DR) yielded approximately tenfold higher colony counts overall and did not show a comparable viability loss for distal edits (Fig. 3G). Instead, reduced viability was confined to a small number of positions, most notably positions 5 and 15.

Across the donor-position series, the predominant outcomes remained the intended HDR allele and the unedited wild-type allele, with non-HDR products representing a minority of reads (Supplementary Fig. 5B). Alternative products for FnCas12a (As DR) were dominated by short deletions centered on the cleavage region, consistent with canonical end joining as a secondary repair outcome (Supplementary Fig. 5B, D-E). In contrast, enAsCas12a accumulated recurrent non-HDR SNPs at later timepoints, which did not resemble a simple canonical NHEJ signature. Together, these data show that both crRNA length and edit position shape Cas12a-mediated precision editing. enAsCas12a supported high HDR across many designs but is more sensitive to distal donor-programmed edits, which are associated with loss of viability and reduced recovery of precise products. FnCas12a (As DR), despite slower editing and stricter crRNA-length requirements, maintained higher viability and more stable editing across distal edit positions.

### Cas12a enables efficient donor-templated editing across diverse genomic loci

Having evaluated enAsCas12a and FnCas12a across different target contexts at the GFP locus, we next asked whether these observations extended to endogenous yeast targets. Because editing efficiency and repair outcome can depend strongly on sequence and chromatin context [16,28,29], we designed crRNAs and donor templates to install SNPs and MNVs (multi-nucleotide variants) at 11 distinct genomic loci (Fig. 4A, Supplementary Table S5). This panel included targets previously reported to span a range of crRNA activities [17,18], allowing us to compare editing regimes across both strong and weak editing contexts. We first focused on enAsCas12a with and without LexA–FHA donor recruitment, because the latter configuration combined rapid editing kinetics with improved viability, two features important for scaling Cas12a editing to pooled libraries. Although editing remained target-dependent, enAsCas12a supported efficient donor-templated repair across a broad range of endogenous loci (Fig. 4B, Supplementary Table S6). As in the GFP assays, targets showed distinct kinetic profiles over the time course (Fig. 4C; Supplementary Fig. 6A). LexA–FHA had a clearer positive effect on HDR at endogenous loci than in the GFP-loss assay (Fig. 4B, Supplementary Fig. 6A, B), increasing average HDR by approximately 10%, although the magnitude remained target dependent and smaller than previously reported for Cas9 [15,26]. LexA–FHA also substantially improved viability, particularly for highly active crRNAs (Fig. 4D; Supplementary Fig. 6C). crRNAs expressed from p*SNR52* produced slightly higher HDR levels than those expressed from p*RPR1* (Fig. 4B, Supplementary Fig. 6 A,B). Non-HDR repair products remained infrequent, with the exception of one target (G10, 13% after 6 generations with p*RPR1*) (Fig. 4E, Supplementary Fig. 7A). The most frequent non-HDR outcome for G10 was an alignment-sensitive +A insertion within an A-rich region adjacent to the intended AT-to-TA edit. Other outcomes included deletions spanning 1, 5, 9 and 29 nucleotides near the cleavage site (Supplementary Fig. 7B, C), consistent with NHEJ [30]. These deletions were reduced to undetectable levels by LexA-FHA (Supplementary Fig. 7). At other targets, detectable non-HDR outcomes were rare and generally below 0.45% (Fig. 4E, Supplementary Fig. 7).

**Figure 4.**
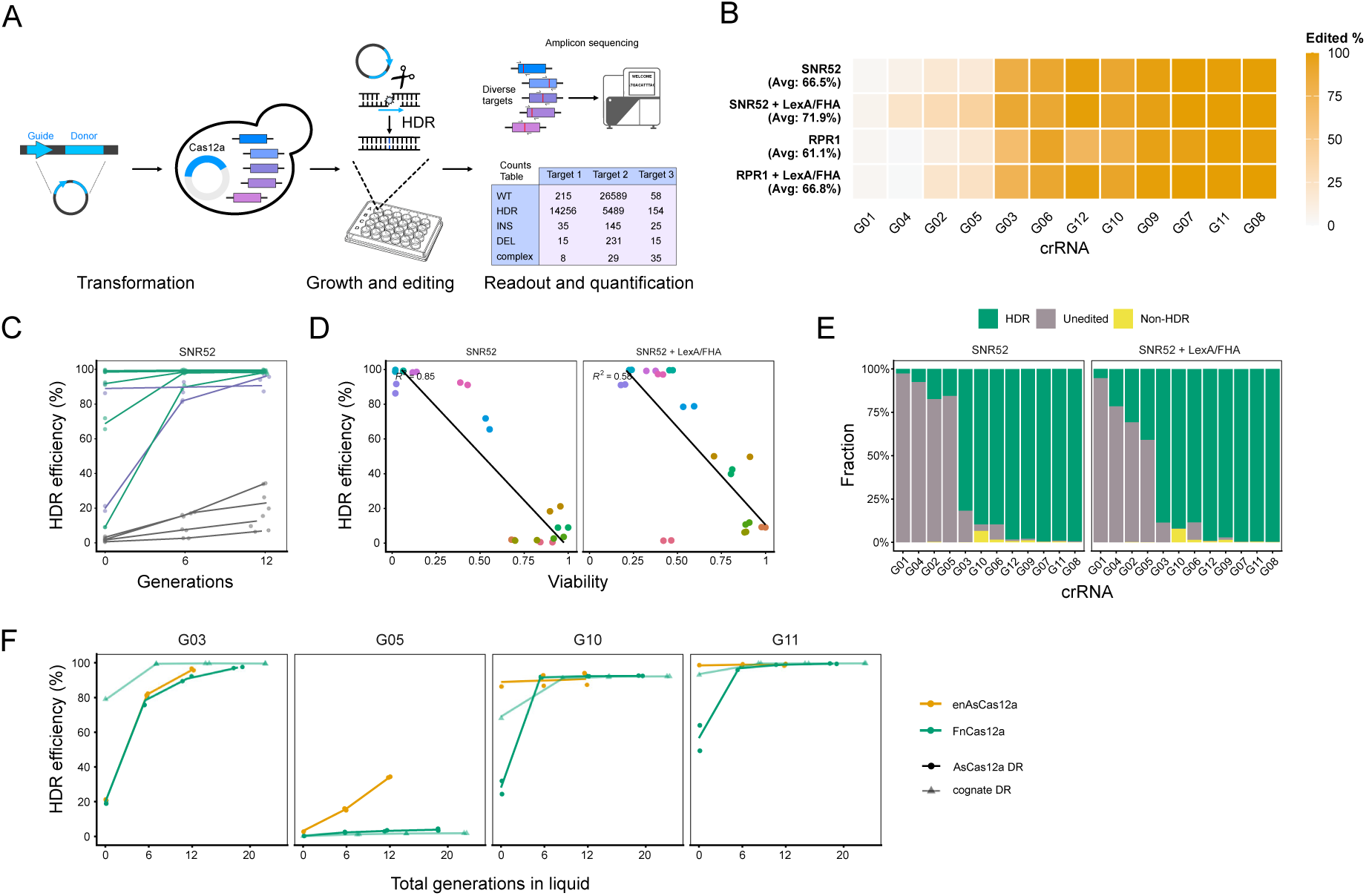
Cas12a supports efficient HDR across endogenous loci with limited non-HDR repair. (A) Assay schematic. Twelve crRNA-donors were designed to introduce single and di-nucleotide edits at endogenous loci. HDR efficiency was quantified by amplicon sequencing after plating and after an additional 6, 12, and 20 generations of editing in liquid culture. Two plated replicates per transformation were propagated independently. (B) HDR efficiency across the endogenous panel after 6 generations of editing in liquid culture. Experiments were performed with p*TEF1*-*enAsCas12a*. Heatmaps show the fraction of reads assigned to the intended HDR allele for each target (columns, ordered by mean HDR efficiency) under the indicated condition (rows), and fill values represent the average of the two replicates. The average HDR efficiency across all targets in each condition is shown on the left. (C) HDR timecourse across the endogenous panel. Points show replicate HDR frequencies and lines replicate means for each crRNA-donor across sampled timepoints. crRNAs are colored by kinetic class as in the GFP-panel: green = high (>90% HDR at 6 generations), grey = low (never exceeds 50% mean HDR), purple = intermediate. (D) Efficiency after plating compared to relative viability. Colored points represent crRNA-donor pairs and lines linear fits within each condition. (E) Editing outcome composition across the endogenous panel after 6 generations of editing in liquid culture. Stacked bars show the fraction of reads classified as HDR, unedited, or non-HDR for each target under the indicated conditions. Targets are ordered by mean HDR efficiency after 6 generations of editing in liquid culture. (F) HDR kinetics at four endogenous targets edited with either enAsCas12a or FnCas12a, with crRNAs expressed from p*SNR52* and without donor recruitment. Symbols show replicate measurements and lines means.

To compare these results with FnCas12a, we tested four targets representing different enAsCas12a behaviors: low activity, slow kinetics, high activity and high activity with detectable non-HDR outcomes. FnCas12a performed poorly with the low-efficiency crRNA G05 regardless of DR identity but reached final HDR levels comparable to or higher than enAsCas12a at the other targets (Fig. 4F, Supplementary Fig. 6D). Consistent with earlier observations, the cognate DR generally accelerated FnCas12a editing relative to the AsDR. FnCas12a did not produce the same NHEJ-associated indels observed for G10 with enAsCas12a, although the recurrent +A insertion was still detected (Supplementary Fig. 7B, C). As with enAsCas12a, low-frequency deletions (<1.5%) near the cleavage site of G03 were suppressed by LexA-FHA.

Finally, we used the GFP-loss and amplicon-sequencing datasets to explore sequence features associated with precision editing. Across panels, total editing and precise HDR were most strongly associated with PAM identity, crRNA GC content, and edit distance from the PAM, although the dominant predictor varied across assays, time points, and nucleases. In the GFP-targeting panel (Fig. 2), which reflects total editing, edit distance explained the largest proportion of variance for enAsCas12a after 6 generations of editing (Δ adjusted R² = 0.376; Supplementary Fig. 8A). Conversely, PAM identity was the dominant predictor for FnCas12a (Δ adjusted R² = 0.787), with TTTC PAMs performing significantly worse than TTTA PAMs (Supplementary Fig. 8A). Lower crRNA GC content was also associated with higher early editing, although this effect was smaller than the PAM-distance and PAM-identity effects (Supplementary Fig. 8A). The same features influenced HDR and total editing at endogenous loci, but with different effect sizes and optima (Supplementary Fig. 8B,C). For enAsCas12a, crRNA GC content was negatively associated with both HDR and total editing at the earliest timepoint (Δ adjusted R² ∼ 0.12), but this effect diminished with additional generations of editing. After six generations, PAM identity explained more variance in both total editing (Δ adjusted R² = 0.105; Supplementary Fig. 8B) and HDR (Δ adjusted R² = 0.088; Supplementary Fig. 8C), with TTTG PAMs performing worse than other motifs. These differences likely reflect the distinct assay formats and targets, uneven representation of design features, and the narrower dynamic range of the GFP panel, where many crRNAs approached maximal editing. More general design rules therefore require broader sampling across crRNAs, donor designs, and genomic loci.

### Genome-wide pooled precision editing with Cas12a

The targeted experiments above established Cas12a as a robust nuclease for donor-templated editing in yeast, but also showed that editing efficiency depends on crRNA, donor, PAM, and target-context features. Defining these dependencies requires measurements across a larger and more diverse set of edits than can be obtained from individual reporter or amplicon assays. At the same time, pooled Cas12a editing would provide an important complement to Cas9-based variant engineering platforms, particularly for variants in T-rich regions or outside convenient Cas9 target space. We therefore developed a proof-of-concept pooled precision editing platform using enAsCas12a with LexA-FHA donor recruitment and a crRNA–donor plasmid architecture analogous to MAGESTIC [15], a high-throughput Cas9-based platform for variant engineering and phenotyping in yeast. We designed 812 editing cassettes to install natural yeast alleles and incorporated genomic barcodes to enable downstream strain identification and pooled phenotyping. The library spanned a broad range of sequence and design features, including GC content, edit position, PAM identity, homopolymer content, and genomic context. Notably, up to 60% of alleles were located in AT-rich regions not proximal to a Cas9 PAM, illustrating the expanded target space enabled by Cas12a. We introduced the pooled library into a yeast strain expressing p*TEF-enAsCas12* and LexA-FHA from a low-copy replicative plasmid. In this format, each transformant carries one crRNA-donor construct, allowing many programmed edits to be generated and propagated in parallel (Fig. 5A).

**Figure 5.**
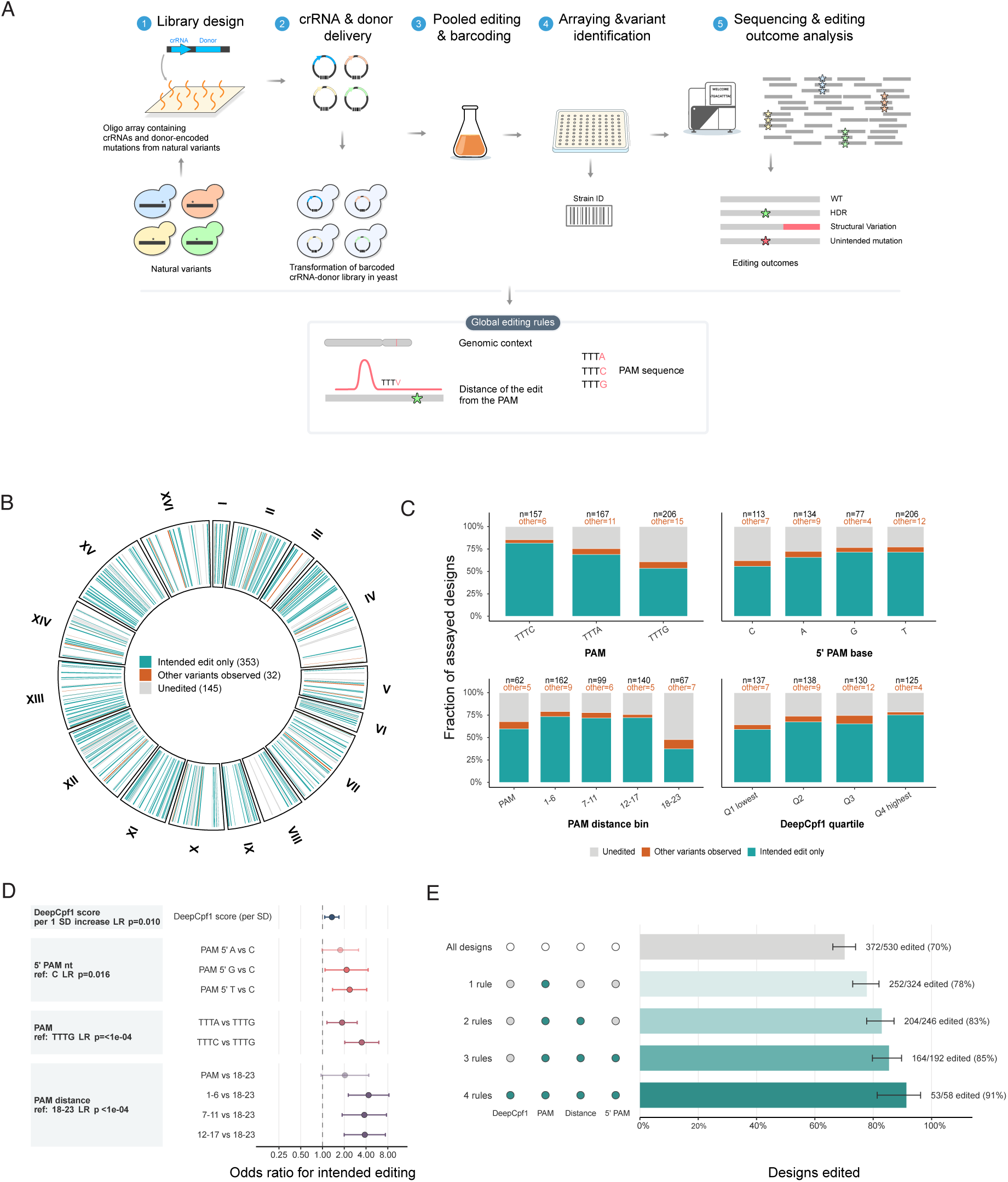
Genome-wide analysis reveals design features associated with successful Cas12a editing of natural genetic variants. (A) Schematic of the pooled editing and editing outcome analysis workflow. Array-synthesized oligonucleotides encoding crRNA-donor cassettes for 812 yeast natural genetic variants were amplified with barcoded primers and subcloned to generate a plasmid library. Yeast cells expressing p*TEF1-enAsCas12a* and LexA-FHA were transformed with the library, edited pools were arrayed to isolate single clones, and editing outcomes were determined by low-coverage whole-genome sequencing. (B) Genome-wide distribution of editing outcomes across the yeast genome. Thin radial bars mark individual assayed designs at their genomic positions and are colored by outcome: intended edit only (teal), unedited (grey), or non-HDR variants (orange). (C) Editing outcome composition across design features. Stacked bars show the fraction of assayed designs (n = 530) classified as correctly edited (teal), unedited (grey), or carrying unintended mutations (orange) across PAM sequences, the nucleotide immediately 5’ of the PAM, PAM-to-edit distance, and DeepCpf1 score quartile. Numbers above bars indicate the number of designs contributing to each feature level. (D) Factors associated with HDR success. Odds ratios were estimated using a multivariable logistic regression model including DeepCpf1 score, PAM sequence, the nucleotide immediately 5’ of the PAM, and PAM-to-edit distance. Points show odds ratios and whiskers indicate 95% confidence intervals. Categorical predictors are shown relative to the indicated reference level; for DeepCpf1 scores, odds ratios indicate the change in odds of successful HDR per 1 SD increase in score. Values above 1 indicate increased odds of successful HDR. (E) Design-feature prioritization. Filled circles indicate favorable included in each subset: DeepCpf1 top quartile, PAM not TTTG, PAM-to-edit distance <18 nt, and 5’ PAM base not C. Horizontal bars show the percentage of edited designs in each subset. Error bars indicate 95% Wilson confidence intervals and labels show the percentage edited and the total number of assayed designs.

Editing outcomes were quantified as described previously [16]. Briefly, edited colonies were arrayed and genotyped by combining recombinase-directed indexing (REDI) [31] for barcode identification with low-coverage whole-genome sequencing using a scalable, extraction-free library preparation method [32]. This workflow recovered strains corresponding to 577 distinct designs (Supplementary Table S7), with some designs represented by multiple independent barcodes. Among 530 designs with at least two reads covering the intended edit site, 70% showed evidence of the programmed edit (Fig. 5B, Supplementary Table S8). Consistent with the smaller-scale assays, non-HDR edits were rare, corresponding to only 6% of the assayed designs. Of these, 25% were near sites previously flagged as high risk for structural variant formation [16] (SCORE cutoff >=0.15). For designs represented by multiple barcodes, editing outcomes were generally concordant across independent isolates (Supplementary Fig. 9A). Because editing was scored as a binary outcome for each recovered clone, detection of an edited isolate indicates that the corresponding cassette can support efficient editing, not that all cells carrying that cassette were edited. Discordant barcode-level outcomes are therefore expected even for active crRNAs, which can still yield unedited clones, although misassignment of low-abundance barcodes may also contribute. Nevertheless, recovery of edited alleles across hundreds of designs shows that enAsCas12a supports efficient pooled donor-templated editing across diverse endogenous targets.

The pooled library also enabled a more systematic analysis of design features associated with editing success. We first tested individual cassette features and found that parameters linked to Cas12a target recognition were the strongest determinants of editing success (Fig. 5C; Supplementary Fig. 9B). PAM-proximal features were strongly associated with editing success. TTTG PAMs performed worse than the other canonical TTTV motifs (logistic regression, *p* = 1.22 x 10^-5^), and the immediately upstream nucleotide also contributed to outcome (*p* = 0.0157), with C being the least favorable base. Edit position relative to the PAM was another significant determinant (*p* = 3.94 x 10^-5^), with edits within the seed-proximal or central spacer region recovered more frequently than distal edits at positions 18-23, consistent with the crRNA length and edit position dependencies observed earlier. In contrast, editing success was not significantly associated with crRNA or donor GC content (*p* = 0.813, *p* = 0.756, Supplementary Fig. 9B).

We next asked whether existing mammalian Cas12a cleavage predictors or RNA-structure features explained HDR recovery. Designs in the highest quartile of enDeepCpf1 scores [12] were enriched for successful editing (global *p* = 0.021, Q4 vs. Q1 *p* = 0.0033), indicating that a mammalian Cas12a cleavage predictor captures part of the signal in our yeast HDR dataset. In contrast, predicted spacer–DR interactions, which could disrupt crRNA folding and have been linked to Cas12a activity [33], were not significantly associated with editing success (Supplementary Fig. 9B). Finally, to identify features that contributed independently, we combined the main predictors in a multivariable logistic regression model. PAM identity, the 5′ PAM-flanking nucleotide, PAM-to-edit distance, and enDeepCpf1 score each retained predictive value, indicating that precision editing success is shaped by multiple partially independent features of target recognition and cleavage (Fig. 5D). The adjusted odds ratios from this model provided a quantitative basis for prioritizing editing cassettes rather than relying on any single feature alone.

Based on these adjusted effects, we derived practical design filters for pooled Cas12a editing. Excluding TTTG PAMs increased the edited fraction from 70.2% to 77.8%. Adding an edit-position filter to exclude distal edits increased this to 82.9%, and further excluding targets with a C immediately upstream of the PAM increased editing to 85.4%. Prioritizing designs in the highest enDeepCpf1 quartile increased the edited fraction to 91.4%, although across a substantially smaller design set (Fig. 5E). The same prioritization criteria also improved performance in the targeted datasets, increasing mean editing from 72.1% to 93.2% among the four retained GFP-targeting crRNAs and from 71.9% to 79.2% among the five retained endogenous targets. Among the four least efficient crRNAs in our targeted genomic panel, two sites were targeted with the unfavorable TTTG PAM, and three had low predicted enDeepCpf1 scores (< 0.5). Thus, these sequence features provide practical criteria for prioritizing cassettes in future pooled Cas12a precision-editing screens.

## Discussion

This study defines key parameters that shape Cas12a-mediated precision editing and establishes a pooled Cas12a platform for genome-scale variant engineering and phenotyping in the powerful model eukaryote *S. cerevisiae*. While previous studies have shown that Cas12a can support HDR in yeast [17–19,21], Cas12a-mediated precision editing has not, to our knowledge, been characterized at this level of mechanistic and design resolution. By systematically varying orthologs, DR configurations, crRNA lengths, expression architectures, donor designs and editing duration, we resolve how these parameters affect not only nuclease activity, but also precise HDR, alternative repair outcomes, and post-edit viability, which are often conflated. This distinction is important because high cleavage activity alone was not sufficient for optimal precision editing and could, in some contexts, reduce recovery of precise edits. We show that precise and viable editing can be improved both by tuning the editing system itself and by prioritizing cassette features that favor successful HDR at genome scale.

Ortholog choice, DR identity and expression architecture generated Cas12a editing regimes with distinct activity, kinetic and viability profiles. Many Cas12a orthologs were highly active in yeast, but maximal activity was not always optimal for precision editing. With their cognate DR, most orthologs supported high maximal levels of overall and HDR editing, yet differed markedly in kinetics and viability. enAsCas12a and FnCas12a were the strongest editors, whereas MbCas12a edited more slowly and AsCas12a showed lower activity under p*TEF1*-driven expression that improved with p*PGK1*-driven expression. crRNA expression architecture provided another layer of control. Pol III promoters generally supported efficient overall and HDR editing, although promoter performance varied with target and nuclease, with p*SNR52* giving the most consistent results across targets. Cas12a ortholog performance therefore depends not only on intrinsic nuclease activity, but also on expression context and editing duration. It is also target dependent, as shown by our broader comparison across endogenous site and consistent with mammalian studies showing target-specific differences in indel formation across Cas12a orthologs [12].

DR identity also shaped Cas12a activity. Although all tested non-cognate Cas12a-DR combinations retained detectable activity, cognate DRs generally improved editing kinetics and maximal efficiency. FnCas12a was especially permissive to the AsCas12a DR, consistent with previous reports of broad DR tolerance for this ortholog [5,18,19]. However, even for FnCas12a, the AsCas12a DR delayed or reduced population-level editing at some targets, indicating that the effect of DR exchange is target dependent and should not be assumed to be universally neutral or uniformly attenuating. LbCas12a and MbCas12a were more strongly impaired, possibly because changes in both loop sequence and loop length make the AsCas12a DR less compatible with these orthologs. This interpretation is consistent with reports that shortened MbCas12a DR loops can compromise activity [34]. DR compatibility should also be considered separately for RNP assembly/cleavage and crRNA precursor processing. FnCas12a tolerated the AsCas12a DR in mature crRNA contexts but was strongly impaired when Pol II-driven expression required processing of a precursor containing non-cognate repeats. In this design, the downstream DR contained a structure-preserving stem base pair change relative to the FnCas12a cognate DR, and similar stem alterations can reduce FnCas12a function in mammalian cells [35]. Impaired precursor processing or reduced mature crRNA production therefore provide plausible explanations for the reduced activity observed here. Overall, these findings show that individual components of the editing machinery, including the Cas12a nuclease, DR, and expression system, can be combined to produce distinct editing regimes, but that each configuration must be evaluated in the relevant context.

Importantly, our data reveal two ways in which highly active editing regimes can reduce recovery of precise HDR products. First, high initial HDR was often followed by a decline in edited cells over time, consistent with continued cleavage after precise repair and secondary loss of HDR products through non-HDR repair, reduced viability, or enrichment of unedited escapers. In contrast, slower or attenuated configurations, such as FnCas12a paired with the AsCas12a DR, often showed gradual accumulation of HDR products and higher viability. Editing duration is therefore an important parameter for maximizing HDR. With highly active configurations, shorter editing may preserve precise HDR products while longer editing benefits slower systems that accumulate edited alleles more gradually. This is particularly relevant for pooled screens, where limiting generations of growth can also reduce spontaneous mutation accumulation and compositional skew. Second, editing activity was strongly coupled to cell viability. Highly active configurations often yielded substantially fewer colonies, whereas attenuated systems, such as FnCas12a paired with the As DR, preserved viability with only modest compromises in editing kinetics or final efficiency. LexA-FHA donor recruitment partially mitigated this trade-off, improving viability across highly active configurations, with the strongest benefit observed for enAsCas12a.

Several mechanistic observations from the targeted assays have direct implications for optimal design. First, crRNA length had nuclease-specific effects. enAsCas12a tolerated substantial shortening and maintained efficient HDR with crRNAs as short as 17 nt, whereas FnCas12a activity dropped sharply below 19 nt. Second, edit position relative to the PAM strongly influenced HDR recovery, particularly for enAsCas12a. We identified local optima in the first bases of the seed region and at positions 7-17, extending previous work that reported maximal editing between positions 12 and 17 of the protospacer [14]. Distal edits were poorly recovered with enAsCas12a and associated with reduced viability, even when the same crRNA was used, showing that donor design can affect editing outcome independently of crRNA activity. One interpretation is that distal edits do not sufficiently disrupt target recognition and therefore permit repeated cleavage after HDR. This is consistent with the crRNA length experiment and with structural and biochemical studies showing that Cas12a target interrogation is directional and that distal protospacer positions contribute differently across orthologs [5,8,9]. Together, these results further underline that Cas12a precision editing is not determined by crRNA activity alone but by the combined effects of crRNA design and expression architecture, donor-encoded edit position, target context, and sampling time, and that these dependencies differ between nucleases. Nuclease choice may therefore be especially important for installing predefined mutations, where the edit position is fixed by the variant of interest and cannot always be moved into the optimal editing window.

The repair outcome data further highlight the suitability of Cas12a for precision editing in yeast. Across assays, the dominant outcomes were the intended HDR allele or retention of the wild-type sequence, whereas non-HDR products were comparatively infrequent. Low-frequency deletions near the predicted cut site were consistent with end-joining repair, in line with expected Cas12a NHEJ outcomes [30,36]. In contrast, recurrent SNPs and +1 insertions did not resemble canonical NHEJ. Because the +1 insertion mapped within a tandem repeat and accumulated over time, it may reflect a sequence-context-associated outcome, such as polymerase slippage, rather than a specific repair pathway. Its persistence could also be favored if it disrupts re-cleavage of the edited locus. Importantly, these alternative products were not the predominant outcome. The main competitor to precise editing was retention of the wild-type allele, indicating that improving edit recovery in yeast will require promoting productive HDR and survival, not only suppressing alternative repair. LexA–FHA donor recruitment addressed these requirements to different extents. It further suppressed some rare non-HDR products, increased HDR modestly and in a target-dependent manner, and resulted in a strong viability benefit. Although the HDR increase was less pronounced than reported for Cas9-based systems, where donor recruitment can strongly enhance HDR for weak guides [15,16,26], the viability benefit is consistent with previous donor-recruitment studies and particularly relevant for pooled editing. By lowering the fitness cost of editing, donor recruitment can limit enrichment of unedited survivors or non-functional designs and help preserve library representation for pooled phenotyping [15,26,37].

The pooled genome-wide experiment expanded the analysis from selected designs to hundreds of endogenous targets across diverse genomic contexts, providing a broader empirical basis for Cas12a precision-editing cassette design. Because alternative repair products were generally rare, editing success in the pooled setting was expected to depend primarily on whether cassettes could promote cleavage, productive HDR, survival and recovery of edited clones. Consistent with this, precision editing success was most strongly associated with features linked to Cas12a target recognition and post-edit mismatch tolerance. TTTG PAMs performed worse than other canonical TTTV motifs, a C immediately upstream of the PAM was unfavorable, and edits placed in the distal protospacer were recovered less frequently. The weaker performance of TTTG PAMs is consistent with mammalian profiling studies of Cas12a-mediated indel formation [10,12], suggesting that some principles of Cas12a target recognition transfer across systems. However, the pooled data also show that predictors developed for mammalian indel formation only partially explain HDR recovery in yeast. enDeepCpf1 scores [12] were informative but only moderately discriminative, as expected for a readout that combines cleavage, donor-templated repair, avoidance of repeated cutting, cell survival, and clone recovery. Prioritizing designs based on simple target recognition and edit distance rules substantially enriched the fraction of edited targets from 70.2% to 85.4%, approaching the 89% pooled editing efficiency reported for Cas9-based yeast platforms [16]. For dense variant libraries, such rules can increase the expected edited fraction in final pools, whereas less optimal designs may still be useful when target space is constrained. Although the quantitative design rules derived here are specific to enAsCas12a in the yeast expression and donor-recruitment architecture tested, the underlying principles are likely to be more broadly informative.

Several limitations remain. The binary clone-level outcome used in the pooled experiment identifies cassettes capable of producing edited isolates but does not directly measure editing frequency within the original population. Thus, recovery of an edited clone indicates that a design can support efficient editing, but unedited clones may still arise from the same cassette. In addition, low-coverage genome sequencing enabled scalable genotyping but limits detection of rare or complex repair outcomes. Repair pathway usage will also differ across systems. Yeast is highly HDR-permissive, whereas many plant and mammalian systems are expected to show higher frequencies of end-joining products after Cas12a cleavage. Nevertheless, this HDR-permissive context is a strength for mechanistic dissection, as it allows effects of nuclease choice, crRNA architecture, expression context, edit position, and donor design on precise repair to be resolved without being masked by dominant NHEJ. Thus, although the quantitative optima defined here may vary across organisms and editing architectures, the underlying parameters are likely to be broadly relevant. Yeast therefore provides a tractable system for defining general Cas12a editing principles, including PAM-dependent activity and edit-position effects, that can guide nuclease choice and cassette design in other systems, even when additional interventions are needed to shift repair toward HDR.

Lastly, by integrating Cas12a into a pooled editing platform and coupling programmed editing to genomic barcoding, we enable high-throughput single-variant engineering and pooled phenotyping with efficiencies comparable to Cas9-based methods in yeast. Cas12a has so far been used only sparsely for precise variant engineering at scale. For example, Xie et al. used an improved LbCas12a system to engineer amino acid substitutions from a 160-plasmid library, but without integrated variant barcoding [21]. Our approach extends this domain-scale mutagenesis toward barcoded, genome-scale single-variant engineering and pooled phenotyping. Used alongside Cas9-based platforms, pooled Cas12a editing should substantially broaden the fraction of natural and engineered variation that can be installed and phenotyped in yeast, enabling more complete genotype-to-phenotype maps.

## Conclusions

Together, our results provide a systematic framework for understanding and improving Cas12a-mediated precision editing. By separating nuclease activity from HDR, alternative repair outcomes, editing kinetics and viability, we show that successful variant installation depends on both tuning the editing machinery and selecting favorable crRNA-donor cassette features. These analyses define practical design rules for Cas12a HDR, including effects of PAM identity, local PAM context, crRNA length and donor-encoded edit position. In yeast, these principles enabled a pooled, barcoded enAsCas12a platform that extends precision editing into T-rich and Cas9-limited sequence space. Used alongside Cas9-based systems, Cas12a should broaden the range of variants that can be functionally tested, supporting more complete genotype-to-phenotype mapping.

## Methods

### Yeast strains and growth conditions

The yeast strain background used in all experiments is a strain compatible with MAGESTIC [15] and REDI [31], and is described in both studies. Briefly, the strain is a derivative of BY (S288c) named DHY214 (*Mat-alpha, his3Δ1, leu2Δ0, ura3Δ0, lys2Δ0*) in which genetic defects have been repaired to improve sporulation *[MKT1(30G), RME1 (INS-308A), TAO3(1493Q)]* and mitochondrial genome stability and function [*CAT5(91M), MIP1(661T), SAL1^+^, HAP1^+^*]. Further modifications include deletion of the *FCY1* ORF and integration of a landing pad containing the *FCY1* ORF flanked by I-SceI recognition sites and homologies for integration of constructs on chromosome II. This strain was used for the endogenous panel and pooled genome-wide editing. Reporter assays were performed in strains carrying a genomic *pTDH3-sfGFP-tADH1* reporter (for GFP loss assays) or a *pTDH3-sfGFP(T105*)-tADH1* PTC reporter (for GFP HDR assays) at the landing pad.

Yeast cultures were grown at 30°C in synthetic complete or dropout media as appropriate. Reporter experiments, including the ortholog and promoter comparisons, were performed in SC-URA-LEU medium. The GFP crRNA panel, endogenous panel, and pooled genome-wide editing experiment were performed in SC-URA-HIS medium.

### Cas12a and crRNA-donor expression plasmids

Mammalian-codon-optimized FnCas12a, AsCas12a and LbCas12a coding sequences were based on Zetsche et al. [5]. *S. cerevisiae* codon-pair optimized FnCas12a (pCSN068), LbCas12a (pCSN067), and enAsCas12a (pRDA_174) were obtained from Addgene from Verwaal *et al.* [18]. Cas12a coding sequences were cloned into low copy *URA3* vectors under control of *TEF1* or *PGK1* promoters and *CYC1* terminators. crRNAs were expressed from yeast Pol III promoters p*SNR52*, p*RPR1*, or p*SCR1*, or from the synthetic Pol II promoter syn1 [24]. The *SCR1* promoter corresponds to chromosome V positions 441,652-442,074 in SGD R64-1-1 and includes 335 bp upstream of the annotated *SCR1* ncRNA plus the first 88 bp of *SCR1/YNCE0024W*. All promoter sequences are deposited Supplementary Table S9. Unless otherwise stated, crRNAs used a 23-nt targeting sequence and either the cognate DR for each Cas12a ortholog or the AsCas12a DR. For syn1-driven crRNA-expression, construct used a DR-spacer-DR architecture followed by a t*CYC1*, with either the As DR configuration TAATTTCTACTCTTGTAGAT-spacer-AAATTTCTCCTCTCGGAGAT or the cognate Fn DR configuration TAATTTCTACTGTTGTAGAT-spacer-TAATTTCTACTGTTGTAGAT. The second DR in the As configuration is an engineered repeat sequence compatible with AsCas12a crRNA multiplexing [11]. crRNAs and donors were subcloned into multicopy 2µ plasmids with a *LEU2* marker for ortholog and promoter comparisons and a *HIS3* marker for all subsequent experiments. For LexA-FHA donor recruitment experiments, crRNA-donor plasmids additionally contained LexA-binding sites, as described previously [15], and the donor recruitment machinery (*pDUT1-LexA-FHA-tADH1*) was expressed from the Cas12a plasmid.

### Yeast transformation and viability measurements

Yeast transformations were adapted from a high-efficiency lithium acetate/polyethylene glycol protocol. Single colonies or dense colony material were grown overnight in selective medium, diluted to OD_600_ 0.15 to 0.2 in YPD and grown for approximately 4 h at 30° C to mid-log phase. Cells were washed and resuspended in 0.1M LiOAc to yield 100 µL competent cells per transformation. For each transformation, 100 ng plasmid DNA in 150 µL water was mixed with 50 µL boiled and chilled ssDNA, 100 µl competent cells and LiOAc/PEG solution containing 600 µL 50% PEG 3350 and 100 µL 1M LiOAc. Reactions were incubated 30 min at room temperature and 30 min at 30°C after LiOAc/PEG addition, heat shocked at 42°C for 14 min, resuspended in 200-250 µL 5 mM CaCl_2_ and plated within 10 min on selective agar. Plates were incubated for 48h at 30°C. After 48 h, transformation plates were photographed and colony count used as the primary viability measurement. Counts were counted from plate images using a custom YOLO object-detection model trained on several hundred manually annotated yeast plate photographs. The model showed high agreement with manual counts for low- and moderate-density plates, which covered the range most relevant for editing-associated viability differences, but tended to underestimate counts at high colony densities. Representative detection examples are shown in Supplementary Figure 10. Colony counts were normalized within each experiment to calculate relative viability. For GFP-panel experiments, counts were normalized to the non-targeting control within each condition. Ortholog comparisons used normalization to the maximum colony count within each Cas12a promoter and GFP assay. For the endogenous panel, counts were also normalized to the maximum colony count within each crRNA promoter and donor recruitment condition.

### GFP-loss and GFP-HDR reporter assays

For GFP-loss assays, yeast carrying the integrated *pTDH3-sfGFP* reporter and the relevant Cas12a expression plasmid were transformed with plasmids containing a crRNA and 122-nt donor designed to introduce a TAG PTC at position 105 of the sfGFP coding sequence (T105*). For GFP-HDR assays, yeast carrying the integrated *pTDH3-sfGFP(T105*)-tADH1* PTC reporter were transformed with plasmids containing a crRNA targeting the PTC-containing sfGFP allele and a corresponding donor designed to restore the wild-type sfGFP coding sequence. Each transformation was split into two replicates before plating, and plated replicates were propagated independently for the rest of the experiment. Colonies were washed from selective plates after 48 h (generation 0, although editing starts immediately after transformation), diluted to OD_600_ 0.01 in 48-well plates containing 400 µL selective medium, and propagated through serial liquid outgrowth until 20 generations of editing in liquid culture, with samples taken at generation 6, 12 and 20. Flow cytometry samples were prepared in flat-bottom 96-well plates by mixing 100 µL diluted cell suspension with 100 µL Attune Focusing Fluid. Cells were measured on an Attune NxT flow cytometer, collecting 25,000 events per sample. Flow cytometry files were processed with custom R scripts using the flowCore, ggcyto, flowWorkspace, and tidyverse packages. GFP-positive and GFP-negative populations were separated by identifying the low-density region between fluorescence peaks. GFP-loss editing efficiency was calculated as the fraction of GFP-negative cells, and GFP HDR efficiency as the fraction of GFP-positive cells.

### Growth-curves

Growth assays were performed on a BioTek Synergy H1 plate reader to distinguish basal Cas12a expression burden from growth effects caused by active GFP targeting. For growth measurements, Cas12a was expressed from a galactose-inducible promoter to precisely time the onset of Cas12a expression and target cleavage. Yeast strains carrying the indicated Cas12a ortholog under a galactose-inducible promoter were transformed with the crRNA/DR plasmids as described above and grown on selective plates for 48 h. Cells were washed from plates, diluted to OD_600_ 0.1 in 2% glucose-containing selective media, grown for 4 to 5 h at 30° C, washed twice, diluted again to OD_600_ 0.1 in 2% galactose-containing selective media, and transferred to 96-well plates for automated OD_600_ growth measurements at 30°C with continuous shaking. Measurements were collected at 10-min intervals, with an empty-plasmid control included as a shared reference. Growth analysis used blank-corrected OD_600_ values, with negative values set to zero. Exponential-phase growth rates were estimated from selected growth windows and area under the curve was calculated by trapezoidal integration of blank-corrected OD_600_ over time.

### Design and cloning of GFP and endogenous panels

For the GFP crRNA-panel experiments, 12 GFP-targeting crRNA-donor pairs were selected to span different PAM sequences, target-strand orientations, crRNA GC contents, positions along the sfGFP ORF, and PAM-to-edit distances (Supplementary Table S2). crRNAs targeted positions between 19 and 55% of the ORF, and each 125-nt donor introduced a single-nucleotide substitution. A non-targeting crRNA with minimal homology to GFP or the yeast genome, paired with a donor sequence for a mutation outside the tested GFP panel, was included as a negative control. For crRNA-length experiments, the A04 GFP target was used, and the crRNA targeting sequence was shortened from 23 nt to 16 nt in one-nucleotide increments while keeping the donor constant, with the programmed edit at position 4 relative to the PAM. For edit-position experiments, the same crRNA was used, but the donor-programmed single base substitution was moved across positions 0 to 23 relative to the PAM, with position 0 corresponding to the V in a TTTV PAM. The crRNA length was kept constant at 23 nt. The position 6 donor plasmid contained an unintended sequence error and was excluded from analysis. For both the crRNA length and edit position experiments, the crRNA was expressed from p*SNR52*.

For the endogenous genomic panel, crRNA-donors were designed to install single- or two-nucleotide substitutions at selected genomic loci. *ADE2*-targeting designs were taken from Swiat *et al.* [17] and crRNAs targeting genomic integration sites were adapted from Verwaal *et al.* [18]. The remaining designs targeted noncoding regions across different chromosomes and a synonymous SNP in *TAT1* to avoid strong phenotypes. All crRNAs were 23 nt, donors 125 nt, and designs avoided BspQI and BbsI sites used during cloning. Donors were centered on the intended edit, and local GC content was preserved. Complete crRNA, donor and amplicon-primer sequences are provided in Supplementary Table S9.

crRNA-donor oligos were synthesized as 200-nt oligonucleotide pools (IDT), with crRNA and donor sequences separated by a BbsI restriction site and flanked by priming sequences used for amplification and containing BspQI restriction sites. Pools were resuspended at 20 ng/µL in nuclease-free water, aliquoted, and stored at -20°C. Oligonucleotide pools were amplified with KAPA HiFi HotStart ReadyMix using eight cycles. The amplified oligonucleotide pool and vector backbone were digested with BspQI, followed by dephosphorylation and 0.6x bead cleanup of the vector backbone. Ligations were incubated overnight at 16°C using high-concentration T4 DNA ligase, 250 ng vector and a 1:2 vector-to-insert molar ratio. Ligation reactions were ethanol precipitated and electroporated into *E. cloni* 10G Supreme cells using 1-mm cuvettes, 10 µF capacitance, 600 Ω resistance, and 1800 V. Cells were recovered in *E. cloni* Recovery Medium and plated on LB agar containing carbenicillin. After overnight incubation, colonies were washed from plates and plasmid pools isolated by Miraprep [38]. In a second cloning step, an internal fragment containing a Pol III terminator, bacterial kanamycin resistance cassette, and *HIS3* marker was amplified with primers containing BbsI sites, digested with BbsI and DpnI, and ligated overnight at 16°C into the BbsI-digested and dephosphorylated plasmid library. Ligation products were ethanol precipitated, electroporated into *E. cloni* 10G Supreme cells, recovered for 1 h in recovery medium and for an additional 3 h in LB, and plated on low-salt LB agar containing carbenicillin and kanamycin. Individual colonies were arrayed in 96-well plates, screened by colony PCR, and identified by Sanger sequencing of the donor region. All plasmids were verified by multiplexed long-read Nanopore sequencing using an in-house Tn5 protocol adapted from Circuit-seq [39].

### Transformation and editing of the GFP and endogenous panel

For targeted panels, the base strain carrying a plasmid expressing Cas12a and, where indicated, the LexA-FHA donor-recruitment machinery was transformed with individually isolated crRNA-donor plasmids. After 48 h of growth on selective plates, plates were photographed for colony counting. Cells were washed from plates with SC-URA-HIS, diluted to OD_600_ 1, and inoculated into 1 mL SC-URA-HIS in 24-well plates at OD_600_ 0.01 for the endogenous panel and 0.5 mL SC-URA-HIS in 48-well plates for the GFP panel. Cultures were sampled and passaged at approximately 6-generation intervals for experiments with the *SNR52* promoter. Experiments using *RPR1* were sampled on a fixed-time schedule of 12 h intervals, which closely matched the generation-based *SNR52* sampling. Plots show the corresponding generations for each promoter. Optical density was monitored using a BioTek Synergy H1 plate reader.

### Targeted amplicon sequencing

Amplicon primers were designed with Primer3 to amplify genomic windows containing the intended edit. Amplicon lengths were constrained to 160 to 230 nt, primer-pair melting temperatures were kept within 3°C, and primer specificity was checked against the *S. cerevisiae* genome by BLASTn, requiring at least three mismatches to potential secondary binding sites. Sequencing libraries were prepared in a two-step PCR protocol with primers containing Illumina sequencing adapters and dual sample indices and were sequenced on an Element Biosciences AVITI platform using 2 x 150-bp paired-end sequencing. Demultiplexed reads were inspected, adapter trimmed and merged with fastp version 0.23.2 [40], and reports were summarized with MultiQC [41]. Merged reads were aligned to the *S. cerevisiae* S288c reference genome (SGD R64-1-1) with bbmap version 38.84 [42] using slow mapping and a minimum overlap fraction of 0.9. Alignments were converted to BAM format, sorted, and indexed with samtools version 1.20, and inspected in IGV. 100,000 reads per sample were subsampled for variant analysis. Variants were called with freebayes version 1.3.2 [43] using 30-bp haplotypes, minimum coverage of 50 reads, alternative-allele frequency of 0.1%, and minimum mapping quality specified with -C 30. VCF files were parsed with VCFParser and custom Python 3.9 code and aggregated across samples using custom R scripts. For each sample and locus, reads were classified as unedited, intended HDR, or non-HDR further classified as SNP, MNP, insertion, deletion, and complex outcomes. Haplotype frequencies were calculated as the haplotype read count divided by total read depth at the locus and normalized to 100%. Total non-WT editing was calculated as the sum of intended HDR and non-HDR outcomes. For indel size and location analyses, insertions and deletions were extracted from non-HDR calls, and indel sizes computed from the reference-to-alternative allele length difference. Frequencies were averaged across replicate transformations for each condition and timepoint. For within-type composition plots, indel-size frequencies were normalized to the total signal for that indel type, and deletion signals were mapped relative to the PAM for deletion coverage plots. Alternative-repair outcome plots for the endogenous panel (Figure 4E, Supplementary Figure 7) show generation 6 as a representative time point, as outcome profiles were similar across the time course.

### Pooled natural-variant editing library design and cloning

The pooled editing experiment was designed as a proof-of-concept platform for enAsCas12a-mediated installation of natural yeast alleles. We designed 812 crRNA-donor cassettes targeting natural variants across the yeast genome. Of these, 20% were sampled from alleles previously installed using a Cas9-based pooled editing method [44], and the remainder from SNPs and short indels in the RM11 natural isolate. Designs spanned diverse PAM identities, 5’ PAM-flanking bases, PAM-to-edit distances, crRNA and donor GC contents, homopolymer runs, predicted Cas12a activity scores, and genomic contexts. We prioritized variants in sequence space where Cas12a expands access relative to Cas9, such as AT-rich regions not proximal to canonical Cas9 PAMs. Library amplification oligos are listed in Supplementary Table S9 and crRNA-donor sequences in Supplementary Table S7. Plasmid library cloning was performed as described above, with several modifications. crRNA-donor oligos were synthesized as 200-nt oligonucleotide pools (Twist), with crRNA and donor sequences separated by a BspQI restriction site and flanked by priming sequences used for amplification and containing BbsI restriction sites. Pools were resuspended at 4 ng/µL in nuclease-free water, aliquoted, and stored at -20°C. The backbone contained BbsI sites adjacent to the p*SNR52* promoter and a downstream homology sequence for barcode integration. Library amplification used 12 cycles, and a barcode was introduced with the reverse primer. In the first cloning step, the amplified oligonucleotide pool and vector backbone were digested with BbsI and ligated overnight at 16°C using 250 ng vector at a 1:1 vector-to-insert molar ratio. The second step fragment was amplified with primers introducing BspQI sites and contained a Pol III terminator, bacterial kanamycin resistance cassette, upstream barcode integration homology, *HIS3* marker, and p*GAL-I-SceI-tCYC1* for barcode integration. Ligation was performed using 500 ng vector. All other steps were as described above. Barcodes were linked to crRNA-donor cassettes via amplicon sequencing, as described previously [15]. The pooled library was transformed into yeast expressing p*TEF1-enAsCas12a* and LexA-FHA from a low copy *URA3* plasmid and plated on SC-URA-HIS plates. Transformation was performed as described above but using 1000 ng plasmid library in 100 µL water and 10 µL boiled carrier DNA at 10 mg/mL, and heat-shock was extended to 17 min at 42° C. After 2 days, cells were washed off plates and propagated in SC-URA-HIS medium for 6 additional generations of editing. Genomic barcode integration at the landing pad was induced in SC-HIS medium containing 2% galactose and 0.15% glucose for 5 generations, followed by washing and growth in SC-HIS containing 2% galactose for 14 generations. Cells were plated on SC + 5FC + 5FOA plates for 2 days to cure the *URA3*-containing Cas12a plasmid and *FCY1*-containing crRNA-donor plasmid, washed and outgrown in YPD for 5 generations and stored as glycerol stocks.

### Recombinase-directed indexing and extraction-free low-coverage whole-genome sequencing

Edited library stocks were diluted to OD_600_ of 1 and plated at a density that yielded approximately 500 to 1000 well-separated colonies per plate. After 2 days of incubation at 30° C, colonies were picked with a Hudson RapidPick Lite automated colony-picking platform into 384 arrays on YPAD plates. After 2 days of incubation at 30°C, plates were rearrayed to 1536-colony format using a Singer Rotor and processed via recombinase-directed indexing (REDI) [31]. Briefly, fresh 1536 arrays of the REDI barcode strains were pinned onto the edited colony arrays and incubated for 24 h at 30°C to allow mating. Mated arrays were transferred to diploid-selection and recombination medium (minimal medium supplemented with URA and LEU), and plates incubated for 72 h at 30°C. After recombination and selection, colonies were washed off 1536 plates and genomic DNA extracted. For each plate, the region containing the editing barcode and the REDI positional barcode was amplified with Illumina sequencing adapters. Amplicons were sequenced on an Element Biosciences AVITI platform in 2x150 paired-end mode and edited strains corresponding to REDI positional barcodes identified using custom scripts. Based on the identified barcodes, 1152 strains representing 577 unique designs were cherry-picked from 384-arrays using the Hudson Rapid Pick, stored as 96-well glycerol stocks, and processed for low-coverage WGS. Low-coverage WGS libraries were prepared using an extraction-free Tn5 tagmentation protocol [32], according to the procedure described in File S1 of the referenced study, and sequenced on an Element Biosciences AVITI platform with 2 × 150-bp paired-end reads to a coverage of 5x.

### Editing outcome analysis

Raw WGS sequencing reads were deduplicated with fastp and mapped with bbmap to a custom genome comprising the base strain genome and separate contigs for the integrated barcode cassettes identified by REDI. Variants were called with freebayes in windows spanning 750 bp upstream and downstream of the intended edit site, which was assigned from the barcode identified by REDI. For each recovered clone, reads covering the programmed edit site were used to classify the clone as intended edit only, other variant observed, or unedited. Designs were considered assayed when at least two reads covered the intended edit site and supported the same outcome. Alternative repair outcomes were called when present at an allele frequency of ≥50% with a minimum read coverage of 4.

### Design-feature and statistical analyses

Unless otherwise stated, downstream analyses and plotting were performed in R using packages ggplot2, patchwork, broom, emmeans, ggeffects, ggbeeswarm, stringr, readr, tidyr, dplyr, and forcats. For targeted panels, linear models using stats:: lm() were used to relate editing outcomes to PAM-to-edit distance, crRNA GC content, and PAM identity. Broader model-comparison outputs evaluated the same features against baseline models that adjusted for the relevant experimental background; nested-model ANOVA/F-tests were performed using stats::anova(), and feature contributions summarized as changes in adjusted R^2^. Numeric feature predictions were generated with ggeffects::ggpredict, and PAM-level estimates with emmeans. For the pooled natural-variant library, editing success was analyzed as a binary outcome. Single-feature logistic regressions tested associations between the presence of intended editing outcomes and crRNA-donor cassette parameters. enDeepCpf1 scores were computed from target sequences using the published enDeepCpf1 model and were also analyzed by quartile. A multivariable logistic regression model combined the main predictors retained from the single-feature analysis. Results were reported as odds ratios with 95% confidence intervals. Practical design filters were evaluated by calculating the fraction of edited designs after excluding unfavorable feature classes.

## Availability of data and materials

The raw sequencing files for datasets generated in this study are available on ENA, accession number PRJEB115113. The datasets supporting the conclusions of this article are included within the article and its additional files. Code to process sequencing and flow cytometry data is deposited on Github: https://github.com/vbstagui98/cas12a_precision_editing.

## Competing interests

The authors declare that they have no competing interests.

## Funding

This work was supported by an FWO junior research project (G012524N) to S.C.V.

## Authors’ contributions

S.C.V, A.D. and V.S.B conceived and designed the study and wrote and edited the manuscript. A.D., V.S.B., J.N. and D.T. performed experiments. A.D. and V.S.B. analyzed data, and S.C.V provided input to the analysis. S.C.V. was responsible for the coordination of the study. All authors read and approved the final manuscript.

## Acknowledgements

Mammalian codon optimized Cas12a constructs were kindly provided by K. Herbst and M. Knop, and were previously described in Buchmuller, Herbst *et al.* 2019 [19]. We are grateful to the VIB Nucleomics Facility for their help in sequencing and the VIB Data Core for providing support on HPC resources used by this study on computing tasks.

---

**Supplementary Figure 1.**
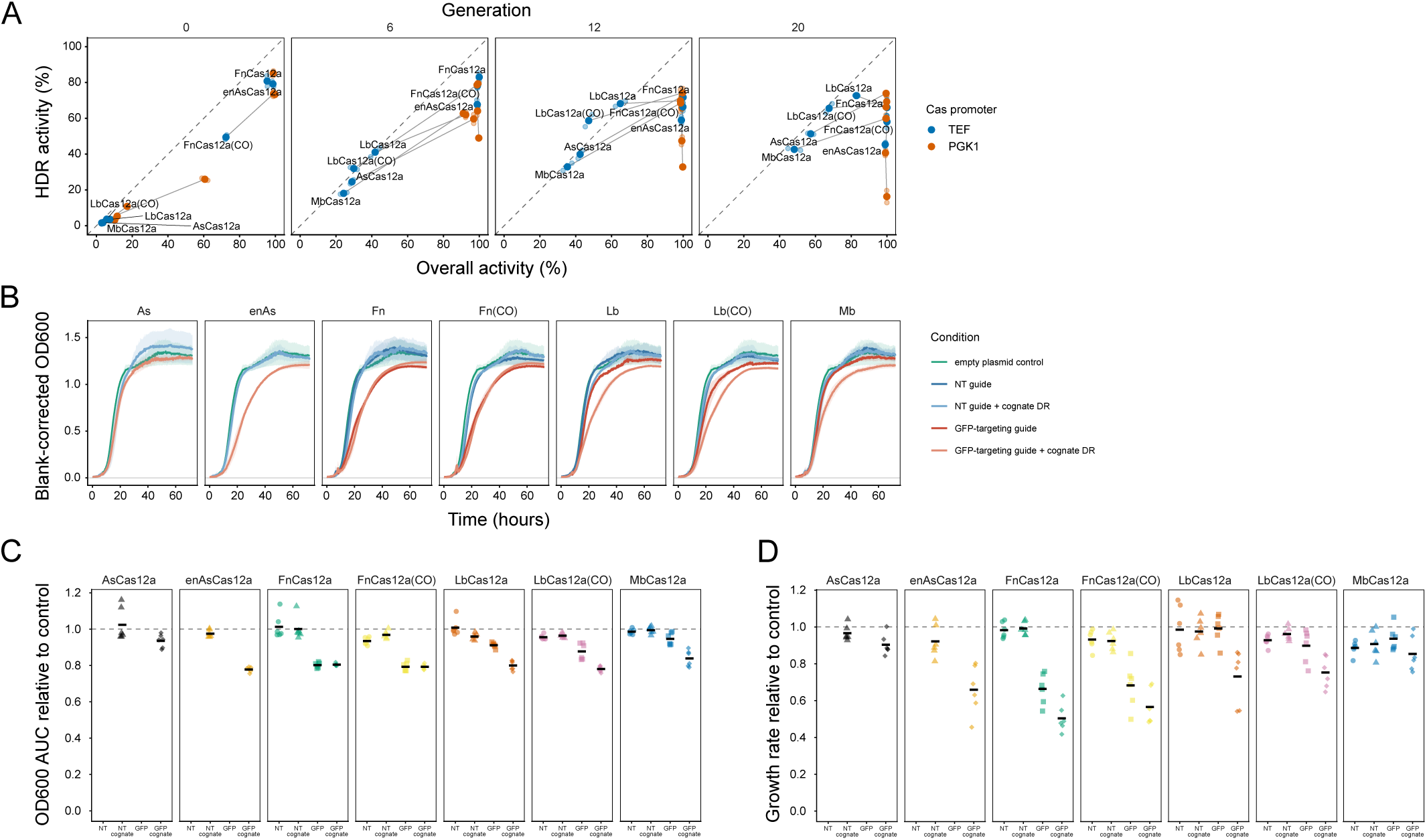
Overall editing activity, HDR activity, and growth effects across Cas12a constructs. (A) Relationship between total editing (GFP loss assay) and HDR editing (GFP restoration assay) across Cas12a orthologs, Cas promoters and timepoints. Colors indicate pTEF- and pPGK1-driven Cas12a expression, and grey lines connect the two promoters for a given ortholog. (B) Blank-corrected OD_600_ growth curves for strains carrying the indicated Cas12a variant and crRNA/DR. Lines show mean OD_600_ over time (n = 6 replicates) and ribbons show 95% confidence intervals. The empty-plasmid control is included as a shared reference. Area under the blank-corrected OD_600_ curve (C) and exponential-phase growth rate (D) for the indicated Cas12a variant and crRNA/DR, normalized to the empty-plasmid control. Points represent individual wells, horizontal bars show condition means, and the dashed line marks the empty-plasmid baseline.

**Supplementary Figure 2.**
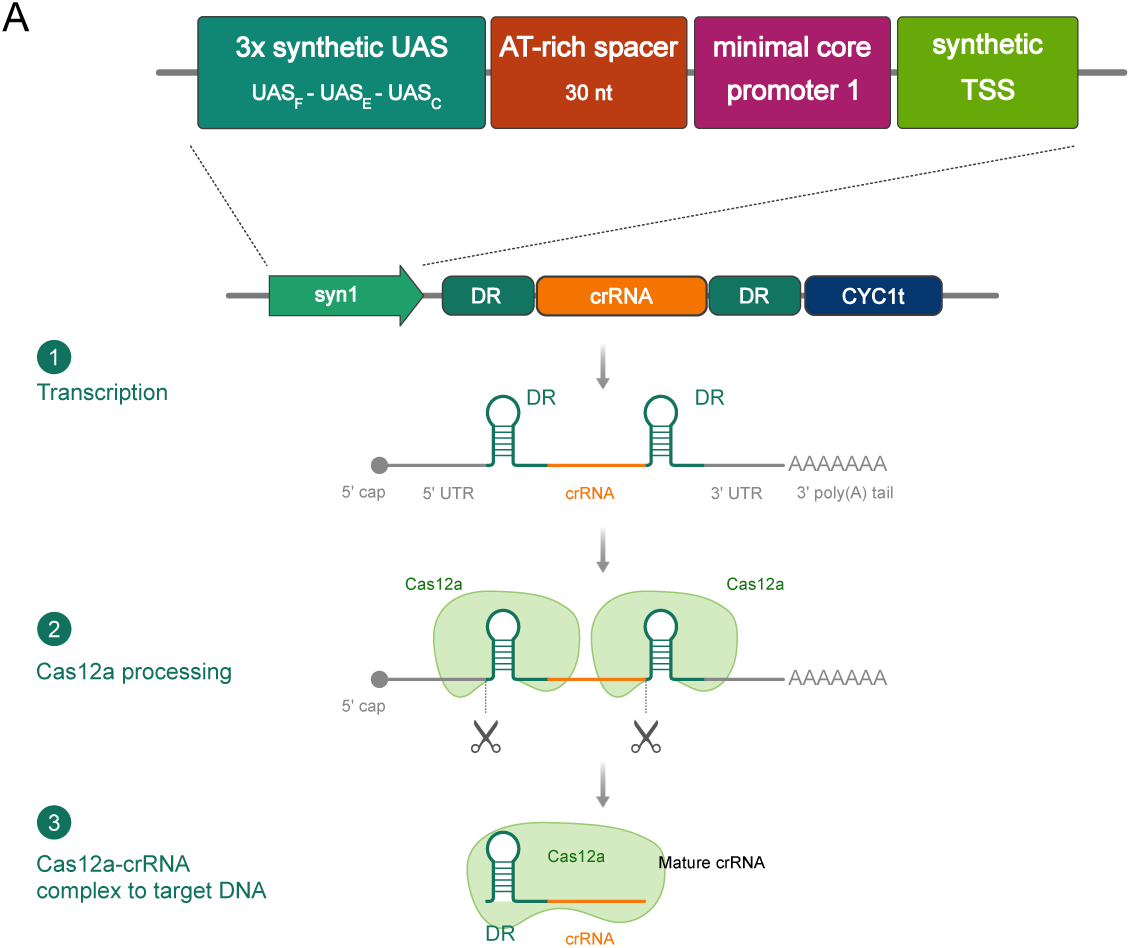
Synthetic Pol II crRNA-expression architecture. (A) Conceptual design of the syn1 Pol II crRNA-expression architecture. The syn1 promoter combines a 3x synthetic UAS, a 30-nt AT-rich spacer, minimal core promoter 1, and a synthetic transcription start site upstream of a DR-crRNA-DR cassette followed by the CYC1 terminator. Transcription produces a capped and polyadenylated RNA containing the crRNA flanked by DRs; Cas12a processes the DRs to release a mature crRNA.

**Supplementary Figure 3.**
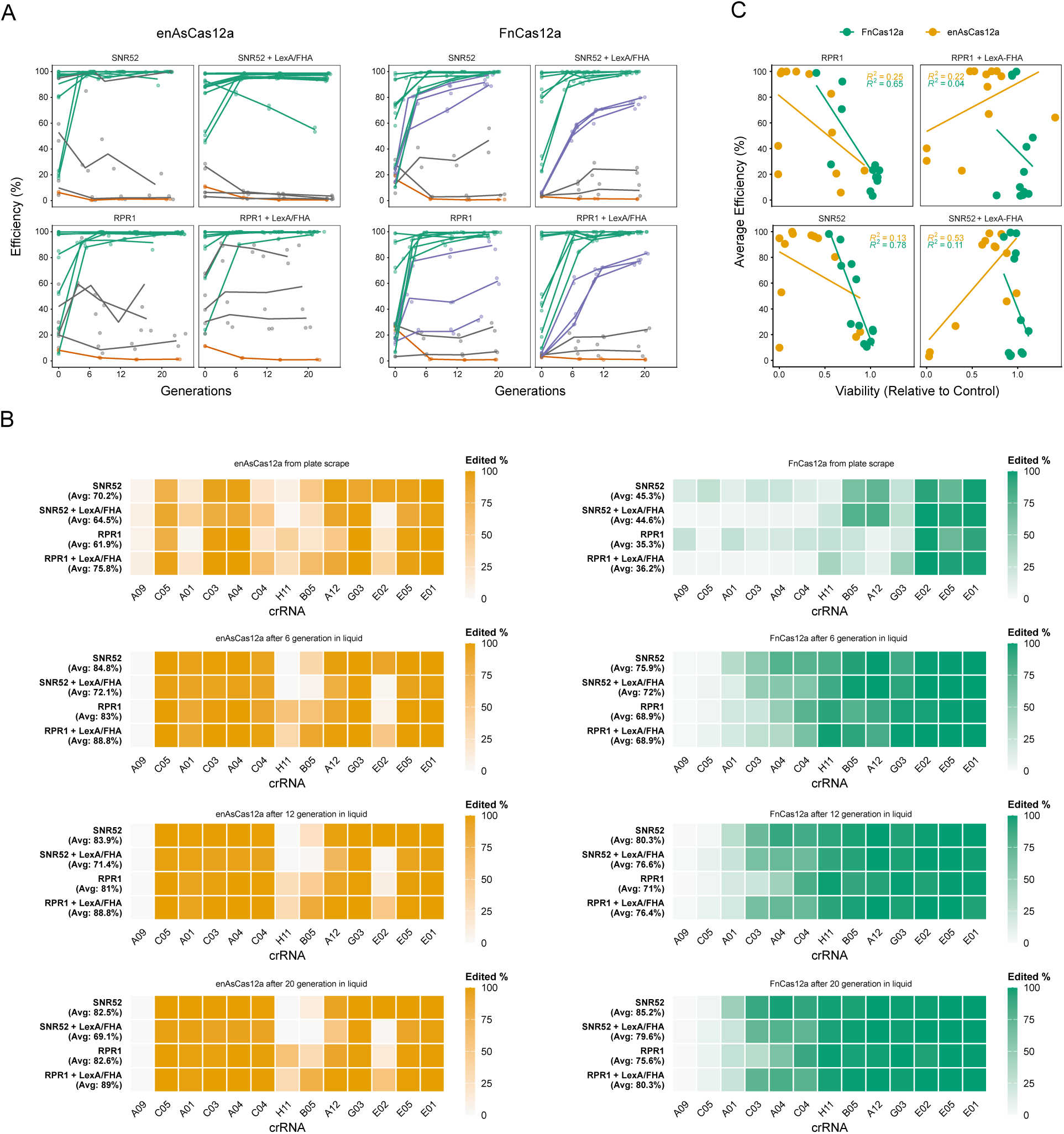
Extended editing time-course analysis in the GFP panel. (A) GFP-loss kinetics across the GFP crRNA panel under the indicated condition. Points show plated replicates and lines replicate means for each crRNA-donor pair. Colors indicate kinetic classes assigned separately for each nuclease: green = high (>90% mean editing at 6 generations), grey = low (never exceeding 50% mean editing), purple = intermediate. (B) Editing efficiency (fraction of GFP-negative cells) across the GFP crRNA panel under the indicated conditions. Heatmaps show average editing efficiencies across replicates. Columns correspond to the non-targeting control followed by GFP-targeting crRNA-donor pairs ordered as in Fig. 2B, and the average editing efficiency of targeting crRNAs in each condition is shown on the left. (C) Editing efficiency compared to relative viability after plating across the GFP crRNA panel. Points show plated replicates and lines ordinary least squares linear regressions. Relative viability was calculated from colony counts normalized to the non-targeting control. In B and C, colors indicate Cas12a ortholog (gold = enAsCas12a, green = FnCas12a (As DR)).

**Supplementary Figure 4.**
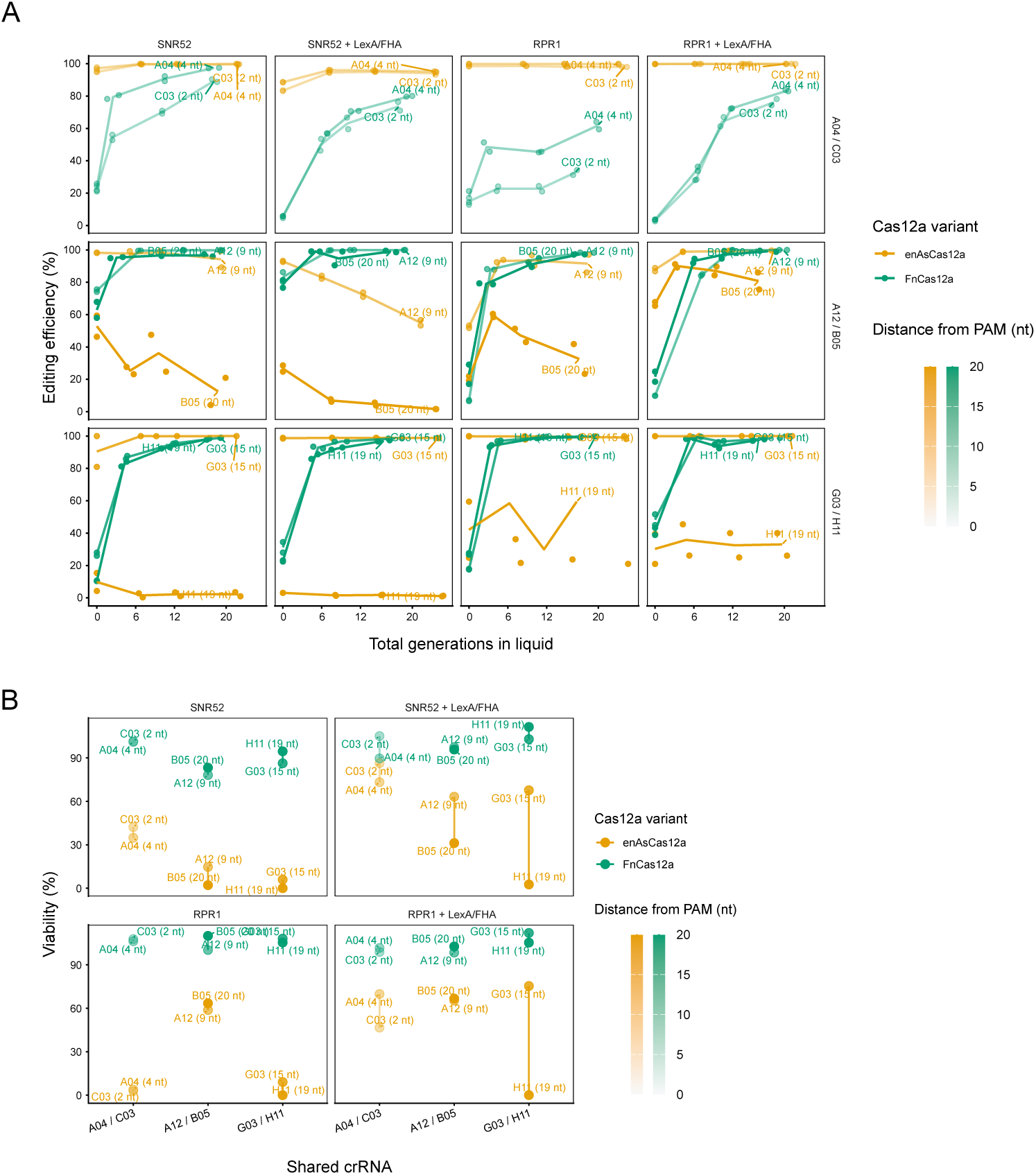
Matched crRNA–donor comparisons reveal donor-position effects independently of crRNA identity. For three crRNAs, donor position was varied by using two donor designs that placed the edit at different distances from the PAM. Labels indicate the well position and PAM distance of each donor design for each pair. Gold points and lines indicated crRNA-matched pairs tested with enAsCas12a; green points and lines indicated matched pairs tested with FnCas12a (As DR); point shading encodes PAM-to-edit distance. (A) Editing kinetics. Rows show shared-crRNA pairs and column show different crRNA-promoter and donor-recruitment conditions. Points indicate replicate measurements, lines show replicate means. (B) Relative viability, calculated as colony counts normalized to the non-targeting control within each Cas12a variant, crRNA promoter and donor recruitment combination.

**Supplementary Figure 5.**
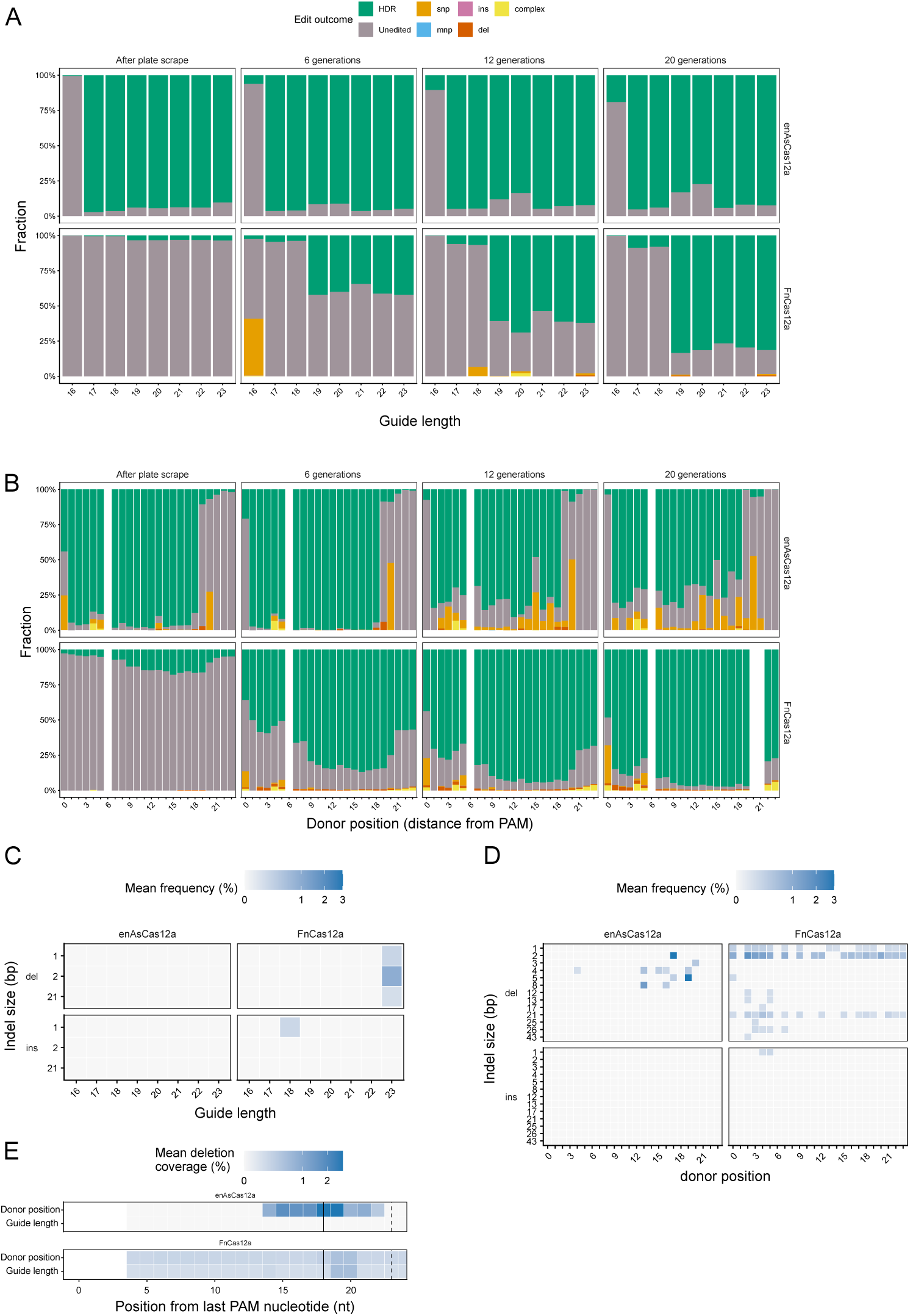
Repair outcomes in crRNA-length and edit-position assays. (A) Repair outcome composition across crRNA lengths (A) and edit positions (B). Bars show the fraction of reads classified as donor-templated HDR, unedited, or alternative outcomes at each sampled timepoint. Indel-size frequency across the crRNA-length (C) and edit-position series (D). Heatmaps show mean non-HDR insertion and deletion frequencies and sizes. (E) Relative deletion coverage across the protospacer. Heatmaps show mean deletion signal at each position across all crRNA lengths (top rows) and edit positions (bottom rows) after 6 (enAsCas12a) or 12 (FnCas12a) generations of editing in liquid culture. Black vertical lines indicate Cas12a staggered breaks.

**Supplementary Figure 6.**
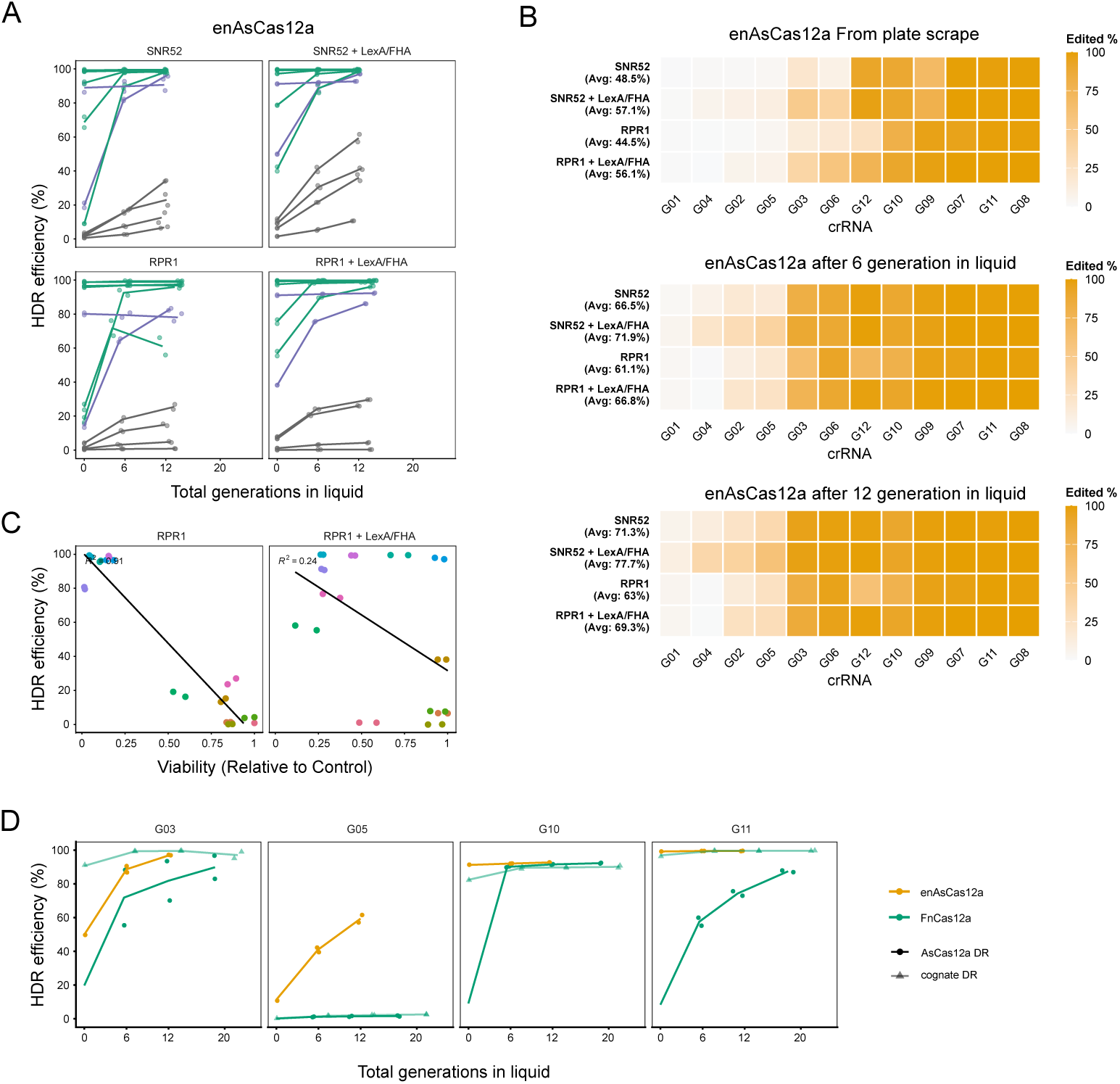
Extended HDR time-course analysis across endogenous targets. (A) HDR kinetics across all sampled timepoints in the indicated conditions. Points show replicate HDR frequencies and lines replicate means for each crRNA-donor pair. (B) Heatmaps show the fraction of reads assigned to the intended HDR allele in the indicated conditions. Columns represent targets ordered by mean HDR efficiency across all conditions, rows indicate experimental conditions, and the average HDR efficiency across all targets in each condition is shown on the left. (C) Efficiency-viability tradeoff. Editing efficiency after plating compared to relative viability, calculated from normalized colony counts (see Methods). Points represent crRNA-donor pairs and lines linear fits. (D) HDR kinetics at targets edited with enAsCas12a or FnCas12a. The plotted condition uses p*TEF*-driven nuclease expression and p*SNR52*-driven crRNA expression with donor recruitment. enAsCas12a trajectories are shown for comparison with FnCas12a paired with either the As DR or its cognate DR. Symbols show replicate measurements and lines show means.

**Supplementary Figure 7.**
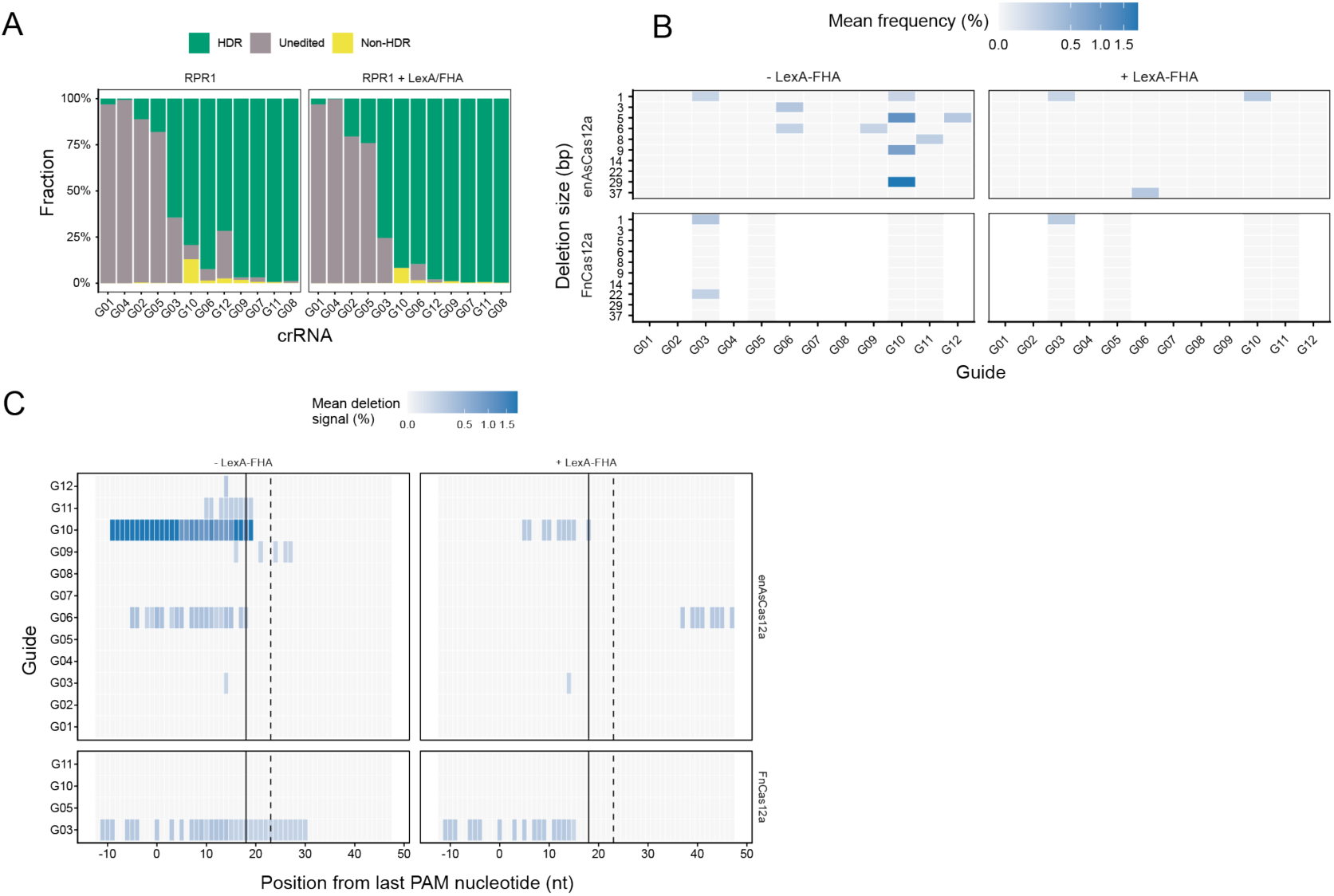
Repair outcomes across endogenous targets. (A) Editing-outcome composition across the genomic target panel. Stacked bars show the fraction of reads classified as intended HDR, unedited, or non-HDR for each target under pRPR1 crRNA expression with or without LexA-FHA. (B) Pattern and frequency of non-HDR deletions at genomic targets. Heatmaps show the mean non-HDR deletion frequencies and deletion sizes after 6 (enAsCas12a) or 12 (FnCas12a) generations of editing in liquid culture averaged across the two crRNA promoters. (C) Relative deletion coverage across the protospacer. Heatmaps show the mean deletion signal at each position for each nuclease and donor recruitment condition after 6 (enAsCas12a) or 12 (FnCas12a) generations of editing in liquid culture averaged across the two crRNA promoters. The black vertical lines mark the sites of the Cas12a staggered break.

**Supplementary Figure 8.**
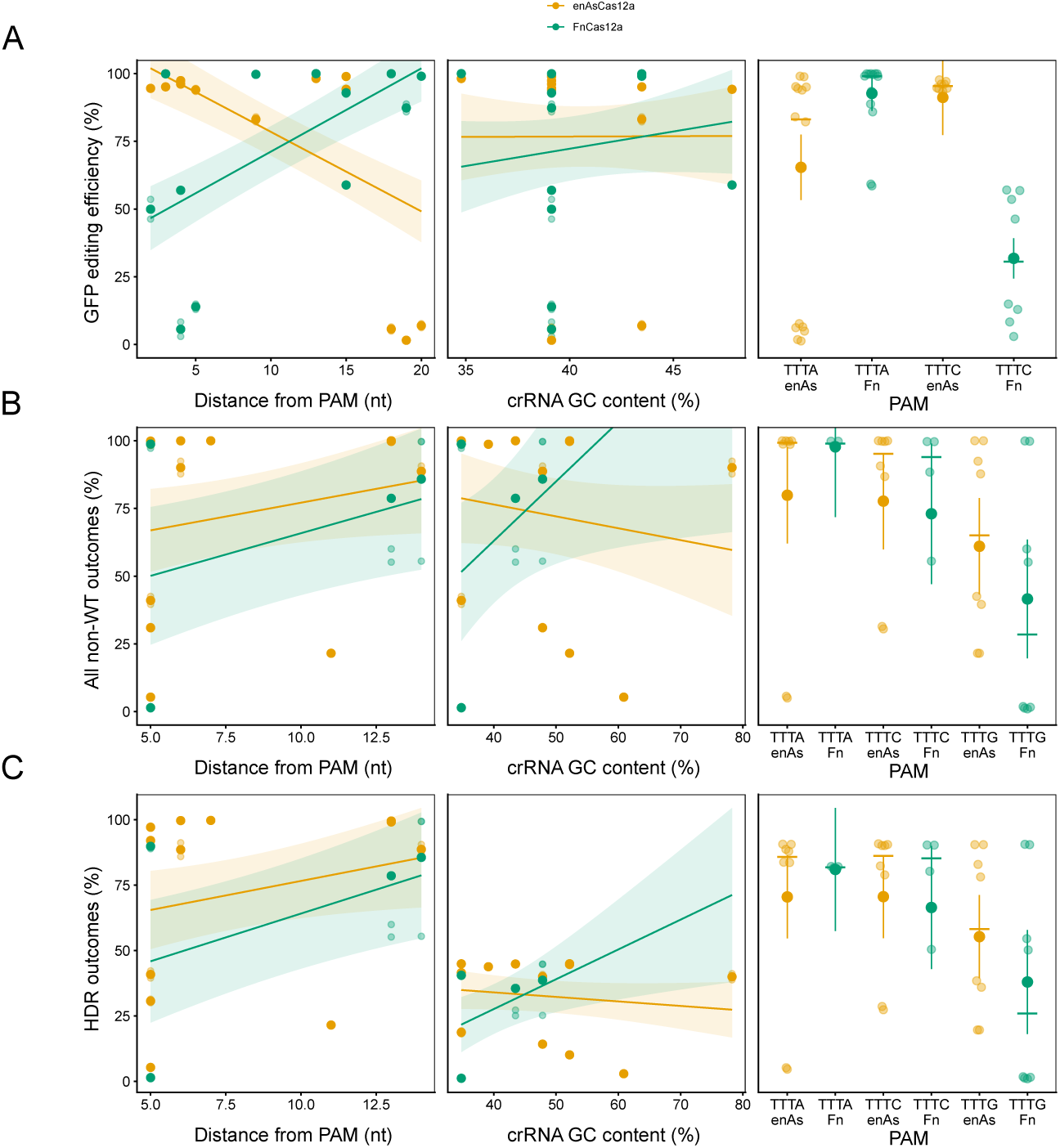
Analysis of design features in the SNR52 + LexA-FHA condition. Efficiency after 6 generations of editing in liquid culture for distinct PAM-to-edit distances, crRNA GC content, and PAM sequences under pSNR52-driven crRNA expression with LexA-FHA donor recruitment. Yellow = enAsCas12a, green = FnCas12a with As DR. (A) Total editing approximated by GFP loss in the GFP panel; (B) Total editing (all non-WT reads) in the endogenous target panel; (C) HDR editing in the endogenous target panel. Smaller semi-transparent points represent replicate measurements and solid points denote mean of the two replicates. Lines and shaded bands show fitted predictions and 95% confidence intervals from variant-specific linear models. For PAM identity, the horizontal bars median values within each PAM group, larger points indicate predicted means and vertical lines 95% confidence intervals from linear models.

**Supplementary Figure 9.**
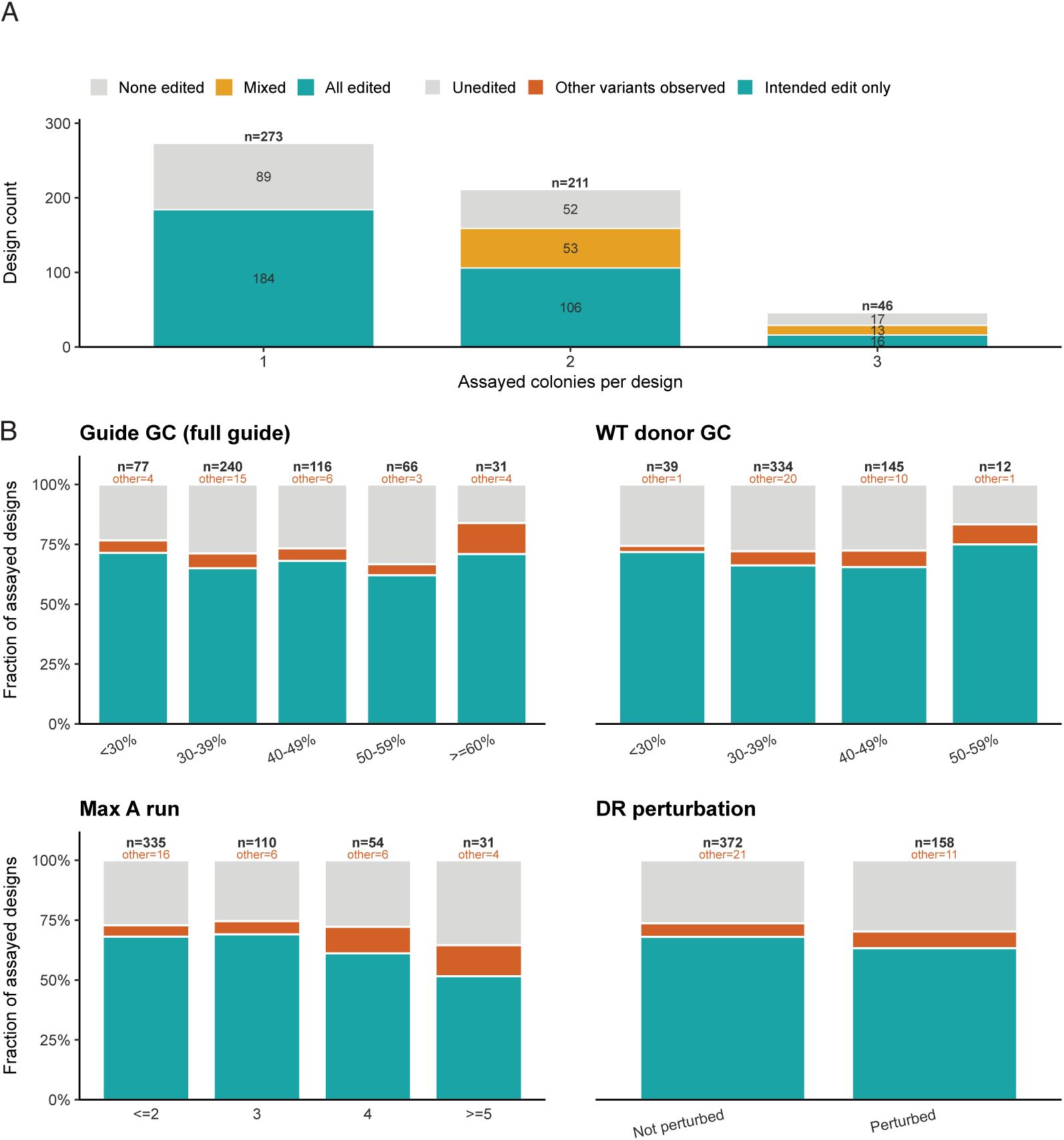
Barcode-level concordance and additional cassette-feature analyses for the pooled Cas12a library. (A) Barcode-level concordance of editing outcomes for designs recovered multiple times. For each design represented by more than one barcode, editing classifications were compared across independently assayed clones. Stacked bars show the number of designs for which all clones were edited (teal), no clones were edited (grey), or barcodes supported mixed outcomes (gold). (B) Fraction of assayed designs classified as intended edit only (teal), other variants observed (orange), or unedited (grey) across design features. Panels stratify designs by crRNA GC content, wild-type donor GC content, maximum A homopolymer run length, and DR perturbation status. Labels report the number of assayed designs in each bin.

**Supplementary Figure 10.**
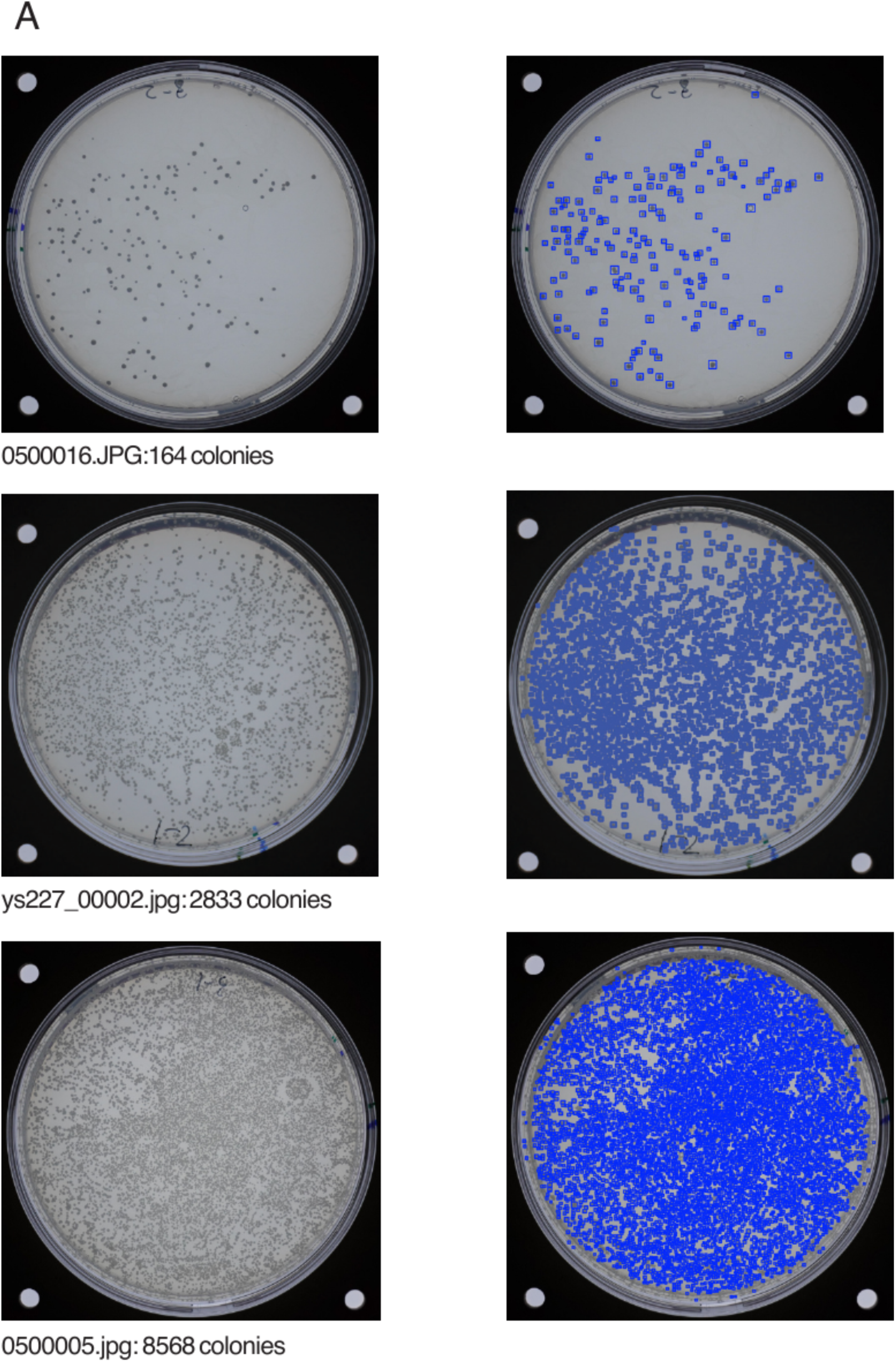
Representative images for the automated colony-counting workflow. (A) Transformation plates with varying colony densities used to measure viability across experiments. For each example, the original image is shown on the left and the processed image on the right, with detected colonies marked by blue rectangles. Source filenames and detected colony counts are indicated below each image pair.

## Notes

### Competing Interest Statement

The authors have declared no competing interest.

## References

1. Jinek M, Chylinski K, Fonfara I, Hauer M, Doudna JA, Charpentier E. A Programmable Dual-RNA-Guided DNA Endonuclease in Adaptive Bacterial Immunity. Science. 2012;337:816–21. 10.1126/science.1225829

2. Xue C, Greene EC. DNA Repair Pathway Choices in CRISPR-Cas9-Mediated Genome Editing. Trends in Genetics. 2021;37:639–56. 10.1016/j.tig.2021.02.008

3. Shalem O, Sanjana NE, Hartenian E, Shi X, Scott DA, Mikkelsen TS, et al. Genome-Scale CRISPR-Cas9 Knockout Screening in Human Cells. Science. 2014;343:84–7. 10.1126/science.1247005

4. Paquet D, Kwart D, Chen A, Sproul A, Jacob S, Teo S, et al. Efficient introduction of specific homozygous and heterozygous mutations using CRISPR/Cas9. Nature. 2016;533:125–9. 10.1038/nature17664

5. Zetsche B, Gootenberg JS, Abudayyeh OO, Slaymaker IM, Makarova KS, Essletzbichler P, et al. Cpf1 Is a Single RNA-Guided Endonuclease of a Class 2 CRISPR-Cas System. Cell. 2015;163:759–71. 10.1016/j.cell.2015.09.038

6. Sudhakar S, Barkau CL, Chilamkurthy R, Barber HM, Pater AA, Moran SD, et al. Binding to the conserved and stably folded guide RNA pseudoknot induces Cas12a conformational changes during ribonucleoprotein assembly. Journal of Biological Chemistry. 2023;299:104700. 10.1016/j.jbc.2023.104700

7. Fonfara I, Richter H, Bratovič M, Le Rhun A, Charpentier E. The CRISPR-associated DNA-cleaving enzyme Cpf1 also processes precursor CRISPR RNA. Nature. 2016;532:517– 21. 10.1038/nature17945

8. Swarts DC, Van Der Oost J, Jinek M. Structural Basis for Guide RNA Processing and Seed-Dependent DNA Targeting by CRISPR-Cas12a. Molecular Cell. 2017;66:221–233.e4. 10.1016/j.molcel.2017.03.016

9. Strohkendl I, Saifuddin FA, Rybarski JR, Finkelstein IJ, Russell R. Kinetic Basis for DNA Target Specificity of CRISPR-Cas12a. Molecular Cell. 2018;71:816–824.e3. 10.1016/j.molcel.2018.06.043

10. Kim D, Kim J, Hur JK, Been KW, Yoon S, Kim J-S. Genome-wide analysis reveals specificities of Cpf1 endonucleases in human cells. Nat Biotechnol. 2016;34:863–8. 10.1038/nbt.3609

11. DeWeirdt PC, Sanson KR, Sangree AK, Hegde M, Hanna RE, Feeley MN, et al. Optimization of AsCas12a for combinatorial genetic screens in human cells. Nat Biotechnol. 2021;39:94–104. 10.1038/s41587-020-0600-6

12. Chen P, Wu Y, Wang H, Liu H, Zhou J, Chen J, et al. Highly parallel profiling of the activities and specificities of Cas12a variants in human cells. Nat Commun. 2025;16:3022. 10.1038/s41467-025-57150-9

13. Richardson CD, Ray GJ, DeWitt MA, Curie GL, Corn JE. Enhancing homology-directed genome editing by catalytically active and inactive CRISPR-Cas9 using asymmetric donor DNA. Nat Biotechnol. 2016;34:339–44. 10.1038/nbt.3481

14. Schubert MS, Thommandru B, Woodley J, Turk R, Yan S, Kurgan G, et al. Optimized design parameters for CRISPR Cas9 and Cas12a homology-directed repair. Sci Rep. 2021;11:19482. 10.1038/s41598-021-98965-y

15. Roy KR, Smith JD, Vonesch SC, Lin G, Tu CS, Lederer AR, et al. Multiplexed precision genome editing with trackable genomic barcodes in yeast. Nat Biotechnol. 2018;36:512–20. 10.1038/nbt.4137

16. Li S, Vonesch SC, Roy KR, Tu CS, Steudle F, Nguyen M, et al. The editable landscape of the yeast genome reveals hotspots of structural variant formation. Sci Adv. 2025;11:eady9875. 10.1126/sciadv.ady9875

17. Świat MA, Dashko S, den Ridder M, Wijsman M, van der Oost J, Daran J-M, et al. FnCpf1: a novel and efficient genome editing tool for Saccharomyces cerevisiae. Nucleic Acids Research. 2017;45:12585–98. 10.1093/nar/gkx1007

18. Verwaal R, Buiting-Wiessenhaan N, Dalhuijsen S, Roubos JA. CRISPR/Cpf1 enables fast and simple genome editing of *Saccharomyces cerevisiae* . Yeast. 2018;35:201–11. 10.1002/yea.3278

19. Buchmuller BC, Herbst K, Meurer M, Kirrmaier D, Sass E, Levy ED, et al. Pooled clone collections by multiplexed CRISPR-Cas12a-assisted gene tagging in yeast. Nature Communications [Internet]. 2019 [cited 2020 Sept 13];10. 10.1038/s41467-019-10816-7

20. Bennis NX, Anderson JP, Kok SMC, Daran J-MG. Expanding the genome editing toolbox of *Saccharomyces cerevisiae* with the endonuclease *Er* Cas12a. FEMS Yeast Research. 2023;23:foad043. 10.1093/femsyr/foad043

21. Xie W, Cai Z, Bao Z. Benchmarking the PAM compatibility of Cas12a variants for high-throughput yeast genetic variant engineering. Nygård Y, editor. Appl Environ Microbiol. 2025;91:e01618–25. 10.1128/aem.01618-25

22. DeWeirdt PC, Sanson KR, Sangree AK, Hegde M, Hanna RE, Feeley MN, et al. Optimization of AsCas12a for combinatorial genetic screens in human cells. Nat Biotechnol. Nature Publishing Group; 2021;39:94–104. 10.1038/s41587-020-0600-6

23. Holland KL, Blancher I, McKesey M, Silas M, Gandhi S, Nickerson A, et al. RNA Polymerase III Promoters Compatible with CRISPR Gene Regulation in Saccharomyces cerevisiae. ACS Synth Biol. American Chemical Society; 2025;14:3387–400. 10.1021/acssynbio.5c00122

24. Redden H, Alper HS. The development and characterization of synthetic minimal yeast promoters. Nat Commun. Nature Publishing Group; 2015;6:7810. 10.1038/ncomms8810

25. Zhao Y, Boeke JD. CRISPR–Cas12a system in fission yeast for multiplex genomic editing and CRISPR interference. Nucleic Acids Research. 2020;48:5788–98. 10.1093/nar/gkaa329

26. Roy KR, Smith JD, Li S, Vonesch SC, Nguyen M, Burnett WT, et al. Dissecting quantitative trait nucleotides by saturation genome editing [Internet]. Genetics; 2024 [cited 2026 Mar 22]. 10.1101/2024.02.02.577784

27. Jansson-Fritzberg L, Chica B, Latrick C, Olland A, Dementiev A, White A, et al. Mechanistic basis for improved activity of Engineered AsCas12a. Commun Biol. 2026;9:565. 10.1038/s42003-026-09799-1

28. Schep R, Brinkman EK, Leemans C, Vergara X, van der Weide RH, Morris B, et al. Impact of chromatin context on Cas9-induced DNA double-strand break repair pathway balance. Mol Cell. 2021;81:2216–2230.e10. 10.1016/j.molcel.2021.03.032

29. Doench JG, Fusi N, Sullender M, Hegde M, Vaimberg EW, Donovan KF, et al. Optimized sgRNA design to maximize activity and minimize off-target effects of CRISPR-Cas9. Nat Biotechnol. 2016;34:184–91. 10.1038/nbt.3437

30. Slaman E, Kottenhagen L, De Martines W, Angenent GC, De Maagd RA. Comparison of Cas12a and Cas9-mediated mutagenesis in tomato cells. Sci Rep. 2024;14:4508. 10.1038/s41598-024-55088-4

31. Smith JD, Schlecht U, Xu W, Suresh S, Horecka J, Proctor MJ, et al. A method for high-throughput production of sequence-verified DNA libraries and strain collections. Molecular Systems Biology. 2017;13:913. 10.15252/msb.20167233

32. Vonesch SC, Li S, Szu Tu C, Hennig BP, Dobrev N, Steinmetz and LM. Fast and inexpensive whole-genome sequencing library preparation from intact yeast cells. Boone C, editor. G3 Genes|Genomes|Genetics. 2021;11:jkaa009. 10.1093/g3journal/jkaa009

33. Creutzburg SCA, Wu WY, Mohanraju P, Swartjes T, Alkan F, Gorodkin J, et al. Good guide, bad guide: spacer sequence-dependent cleavage efficiency of Cas12a. Nucleic Acids Research. 2020;48:3228–43. 10.1093/nar/gkz1240

34. Liu S, He Y, Fan T, Zhu M, Qi C, Ma Y, et al. PAM-relaxed and temperature-tolerant CRISPR-Mb3Cas12a single transcript unit systems for efficient singular and multiplexed genome editing in rice, maize, and tomato. Plant Biotechnology Journal. 2025;23:156–73. 10.1111/pbi.14486

35. Lin L, He X, Zhao T, Gu L, Liu Y, Liu X, et al. Engineering the Direct Repeat Sequence of crRNA for Optimization of FnCpf1-Mediated Genome Editing in Human Cells. Molecular Therapy. 2018;26:2650–7. 10.1016/j.ymthe.2018.08.021

36. Chen R-D, Yang Y, Liu K-M, Hu J-Z, Feng Y-L, Yang C-Y, et al. Post-cleavage target residence determines asymmetry in non-homologous end joining of Cas12a-induced DNA double strand breaks. Genome Biol. 2025;26:96. 10.1186/s13059-025-03567-w

37. Băţăgui VS, Delhaye A, Vonesch SC. From edits to insights: precision microbial engineering for systems biology. Current Opinion in Microbiology. 2026;89:102705. 10.1016/j.mib.2025.102705

38. Pronobis MI, Deuitch N, Peifer M. The Miraprep: A Protocol that Uses a Miniprep Kit and Provides Maxiprep Yields. Hayes F, editor. PLoS ONE. 2016;11:e0160509. 10.1371/journal.pone.0160509

39. Emiliani FE, Hsu I, McKenna A. Multiplexed Assembly and Annotation of Synthetic Biology Constructs Using Long-Read Nanopore Sequencing. ACS Synth Biol. 2022;11:2238–46. 10.1021/acssynbio.2c00126

40. Chen S, Zhou Y, Chen Y, Gu J. fastp: an ultra-fast all-in-one FASTQ preprocessor. Bioinformatics. 2018;34:i884–90. 10.1093/bioinformatics/bty560

41. Ewels P, Magnusson M, Lundin S, Käller M. MultiQC: summarize analysis results for multiple tools and samples in a single report. Bioinformatics. 2016;32:3047–8. 10.1093/bioinformatics/btw354

42. Bushnell B. BBMap: A Fast, Accurate, Splice-Aware Aligner. Walnut Creek, CA; 2014. https://bbmap.org

43. Garrison E, Marth G. Haplotype-based variant detection from short-read sequencing [Internet]. arXiv; 2012 [cited 2026 June 17]. 10.48550/ARXIV.1207.3907

44. Ang RML, Chen S-AA, Kern AF, Xie Y, Fraser HB. Widespread epistasis among beneficial genetic variants revealed by high-throughput genome editing. Cell Genomics. 2023;3:100260. 10.1016/j.xgen.2023.100260

